# Microfluidic antibody profiling after repeated SARS-CoV-2 vaccination links antibody affinity and concentration to impaired immunity and variant escape in patients on anti-CD-20 therapy

**DOI:** 10.1101/2023.10.22.563481

**Authors:** Ashley Priddey, Michael Xin Hua Chen-Xu, Daniel James Cooper, Serena MacMillan, Georg Meisl, Catherine K Xu, Myra Hosmillo, Ian G. Goodfellow, Rafael Kollyfas, Rainer Doffinger, John R Bradley, Irina I Mohorianu, Rachel Jones, Tuomas P.J. Knowles, Rona Smith, V Kosmoliaptsis

## Abstract

**Background:** Patients with autoimmune/inflammatory conditions on anti-CD20 therapies, such as Rituximab, have suboptimal humoral responses to vaccination and are vulnerable to poorer clinical outcomes following SARS-CoV-2 infection. We aimed to examine how the fundamental parameters of antibody responses, namely affinity and concentration, shape the quality of humoral immunity after vaccination in these patients.

**Methods:** We performed in depth antibody characterisation in sera collected four to six weeks after each of three vaccine doses to wild-type (WT) SARS-CoV-2 in Rituximab-treated primary vasculitis patients (n=14) using Luminex and pseudovirus neutralisation assays, whereas a novel microfluidic-based immunoassay was used to quantify polyclonal antibody affinity and concentration against both WT and Omicron (B.1.1.529) variants. Comparative antibody profiling was performed at equivalent time points in healthy individuals after three antigenic exposures to WT SARS-CoV-2 (one infection and two vaccinations; n=15) and in convalescent patients after WT SARS-CoV-2 infection (n=30).

**Results:** Rituximab-treated patients had lower antibody levels and neutralisation titres against both WT and Omicron SARS-CoV-2 variants compared to healthy individuals. Neutralisation capacity was weaker against Omicron versus WT both in Rituximab-treated patients and in healthy individuals. In the Rituximab cohort, this was driven by lower antibody affinity against Omicron versus WT (median [range] K_D_: 21.6 [9.7-38.8] nM vs 4.6 [2.3-44.8] nM, p=0.0004). By contrast, healthy individuals with hybrid immunity produced a broader antibody response, a subset of which recognised Omicron with higher affinity than antibodies in Rituximab-treated patients (median [range] K_D_: 1.05 [0.45-1.84] nM vs 20.25 [13.2-38.8] nM, p=0.0002), underpinning the stronger serum neutralisation capacity against Omicron in the former group. Rituximab-treated patients had similar anti-WT antibody levels and neutralisation titres to unvaccinated convalescent individuals, despite two more exposures to SARS-CoV-2 antigen. Temporal profiling of the antibody response showed evidence of affinity maturation in healthy convalescent patients after a single SARS-CoV-2 infection which was not observed in Rituximab-treated patients, despite repeated vaccination.

**Discussion:** Our results enrich previous observations of impaired humoral immune responses to SARS-CoV-2 in Rituximab-treated patients and highlight the significance of quantitative assessment of serum antibody affinity and concentration in monitoring anti-viral immunity, viral escape, and the evolution of the humoral response.

## Introduction

SARS-CoV-2 infection (COVID-19) is of ongoing clinical concern for patients with primary systemic vasculitis, particularly those receiving repeated dosing with B cell depleting therapies, such as the anti-CD20 agent, Rituximab. Patients with autoimmune/inflammatory conditions on immunosuppressive therapy, particularly those on anti-CD20 therapies(1, 2), are vulnerable to poorer clinical outcomes following SARS-CoV-2 infection, including hospitalisation and death(3, 4). Given this, and the suboptimal humoral immune responses after SARS-CoV-2 vaccination observed in those receiving anti-CD20 therapies(5–7), subsequent SARS-CoV-2 vaccine doses i.e., “boosters”, have been recommended for these patient groups in several jurisdictions(8–10), including those with primary systemic vasculitis(11).

Humoral immune responses after primary vaccination using vaccines targeted against the original (ancestral, wild-type) strain of SARS-CoV-2 e.g., the mRNA vaccines Pfizer BioNTech BNT162b2 (Pfizer) and mRNA-1273 (Moderna) or the adenovirus-vector based vaccines Oxford-AstraZeneca ChAdOx1 nCoV-19 (AZ) or Ad26.COV2-S (Johnson&Johnson), are often suboptimal among patients receiving anti-CD20 therapies. Specifically, titres of antibodies directed against the spike protein subunit (anti-spike IgG) and/or receptor binding domain (anti-RBD IgG) of SARS-CoV-2 and the proportion of patients on anti-CD20 therapy who seroconvert following primary vaccination are lower compared to those not on such therapy and healthy controls(5, 6, 12, 13). Furthermore, of patients on anti-CD20 therapies who seroconvert after primary vaccination, many have lower neutralisation titres compared to those on other immunosuppressants and healthy controls(14–16). Although neutralising antibody titres and, to a lesser extent, anti-spike IgG and anti-RBD IgG titres, derived from primary vaccination with vaccines targeting the original strain of SARS-CoV-2 correlate with protection against symptomatic infection from the ancestral virus(17), significant reductions in the neutralisation capacity of these antibodies have been observed against subsequent SARS-CoV-2 variants of concern, such the B.1.617.2 variant (Delta)(17) and the B.1.1.529 variant and its sublineages (Omicron)(18–20), which harbour mutations in the spike protein that modify the critical domain for virus-neutralising antibodies(21, 22). SARS-CoV-2 specific T cell responses following vaccination, which may be protective against severe infection(23, 24), appear largely preserved among anti-CD20 treated patients(5) although the clinical significance of these responses in such patients remains unclear. Consequently, primary systemic vasculitis patients receiving anti-CD20 therapies may still be vulnerable to severe SARS-CoV-2 infection even if they develop antibodies following primary vaccination, particularly given the emergence of variants with humoral immune escape properties.

We have recently developed a novel immunoassay, microfluidic antibody affinity profiling (MAAP), for in solution quantification of the fundamental parameters of the antibody response, namely affinity and concentration, directly in serum(25) and used it to characterise antibody profiles against wildtype (WT) SARS-CoV-2 in convalescent sera(26) and to study the role of cross-reactivity as a consequence of memory reactivation after acute SARS-CoV-2 infection(27). MAAP can distinguish samples containing low levels of high-affinity antibodies from samples with high levels of low-affinity antibodies, which would otherwise exhibit the same EC_50_ (half-maximal effective concentration) using an ELISA-based technique(26, 27). Thus, MAAP may allow for a more granular assessment of antibody responses following antigen exposure and provide insights into the potency of the humoral response against emerging variants of concern. Nevertheless, a recent study exploring antibody profiles in pre-Omicron sera from largely immunocompetent patients with a variety of antigenic exposures to SARS-CoV-2 antigens (including after one or more doses of WT vaccine or after infection or both) showed similar antibody affinities against Omicron versus WT or Delta Spike antigens, albeit the timing of serum sampling for antibody assessment in this study was widely variable(28).

Several recently published studies have assessed humoral responses following booster vaccination(s) among cohorts of patients with autoimmune/inflammatory conditions on anti-CD20 therapies, which have included vasculitis patients(29–34), although few have examined primary systemic vasculitis patients specifically(35–37). The primary aim of this study was to perform in depth serological characterisation of antibody responses at pre-specified time points after repeated vaccination (specifically after second and third doses) against SARS-CoV-2 in pre-Omicron sera from Rituximab treated patients with primary systemic vasculitis. Along with serological profiling using solid-phase and neutralisation assays, we capitalised on our recently described microfluidics-based immunoassay to quantify antibody affinity and concentration against wild-type (WT) versus Omicron strains of SARS-CoV-2. We subsequently performed antibody profiling at equivalent time points in a cohort of healthy individuals after similar, repeated antigenic exposure to SARS-CoV-2. Finally, we compared the serological response to repeated vaccination in Rituximab treated patients to that in unvaccinated, convalescent patients after a single exposure to WT SARS-CoV-2 infection. We found suboptimal humoral immunity even after repeated vaccination in Rituximab treated patients and highlight the role of quantitative assessment of serum antibody affinity and concentration in monitoring anti-viral immunity, viral escape, and the evolution of the humoral response.

## Methods

### Cohort Description

Patients with primary systemic vasculitides were recruited from a prospective observational cohort study investigating SARS-CoV-2 vaccine responses among renal populations at the Department of Nephrology at Cambridge University Hospitals NHS Foundation Trust (CUH) (ethics reference: 20/EM/0180), as previously described (38). Patients receiving IVIg or plasma exchange were excluded as potential confounders of vaccine responses. Samples from the health care worker cohort were part of the asymptomatic staff screening programme at CUH, as previously described(39). All staff members were invited to participate in the serological screening programme and provided written informed consent (NIHR BioResource - COVID-19 Research cohort; ethics reference: 17/EE/0025, IRAS ID: 220277). The study participants in the convalescent patient cohort were recruited between March 2020 and July 2020 from patients attending CUH with nucleic acid confirmed diagnosis of COVID-19 (ethics reference: 17/EE/0025), as previously described(40). All participants provided informed consent.

Blood samples were taken at approximately 4 to 6 weeks after each sensitisation/vaccination event (1A, 2A or 3A) with flexibility to align with routine clinical tests and access to blood testing facilities wherever possible. Blood samples for the convalescent cohort were taken at the time points specified (1A: one month and 1B: three months from diagnosis of COVID-19). Clinical data collected from electronic medical records and patient interviews included baseline demographics, changes to immunosuppressive medication over time, and data on episodes of SARS-CoV-2 infection.

### Fluorescent labelling of proteins

Recombinant SARS-CoV-2 Spike RBD proteins from WT (#40592-V08H) and Omicron (B.1.1.529; #40592-V08H121) strains were purchased from Sino Biological and labelled with AlexaFluor 647 as previously described(25). In brief, 100μg SARS-CoV-2 RBD protein was re-suspended in H_2_O and mixed with NaHCO_3_ (pH 8.3) buffer to a final concentration of 100mM. DMSO-reconstituted AlexaFluor™ 647 N-hydroxysuccinimidyl ester (ThermoFisher) was added at a 3-fold molar excess and the reaction was incubated at room temperature for 1h. The reaction mixture underwent size-exclusion chromatography in PBS, pH 7.4 using an ÄKTA pure system and a Superdex 200 Increase 10/300 column (Cytiva) to separate the labelled protein from the free fluorophore. Labelled proteins fractions were pooled, concentrated using an Amicon Ultra-0.5 10K centrifugal filter device (Millipore) and glycerol was added to a concentration of 10% (w/v) prior to snap freezing in LN_2_ and storage at -80°C. Upon thawing and before use, protein concentrations were quantified via Nanodrop using the molar extinction coefficient.

### Antibody affinity and concentration measurements

The affinity and concentration of anti-RBD antibodies within patient sera were determined using Microfluidic antibody affinity profiling (MAAP)(26). The hydrodynamic radius (R_H_) of 1nM-500 nM AlexaFluor™ 647-labelled Spike RBD proteins were measured in the presence of MAAP buffer (PBS containing 5% human serum albumin (sigma), 10% (w/v) glycerol and 0.05% Tween-20) and varying concentrations (0-50%) of patient serum via microfluidic diffusional sizing (MDS) using the Fluidity One-W or One-M instruments (Fluidic Analytic Ltd.) after a one-hour incubation on ice. The background fluorescence within each diffused and non-diffused microfluidic stream was subtracted from the MDS data for the specific serum concentrations and Bayesian inference analysis was used to constrain the values of affinity (K_D_) and antibody binding sites ([Ab]), as previously described(25, 26). Serum samples that enabled constrained K_D_ and [Ab] values to be measured (with both upper and lower 95% confidence intervals) were considered quantifiable. Those that had an [Ab]/K_D_ ratio of >2 were labelled as fully quantifiable (Q), and those that had a [Ab]/K_D_ of 1-2 were considered to be at the border of sensitivity for full quantification (B; borderline). Samples which yielded no measured increase in the RBD hydrodynamic radius after serum incubation (N; non-binders) and/or those samples that yielded incomplete K_D_ and/or [Ab] bounds (U; unquantifiable due to inability to fully constrain 95% confidence interval lower bounds for both parameters) were deemed non-quantifiable and excluded from subsequent analysis.

### Luminex assay

Serum antibody reactivity to SARS-CoV-2 WT Spike, RBD and nucleocapsid proteins was assessed using a UKAS accredited Luminex platform, as previously reported(41). In brief, patient serum diluted 1/100 was incubated for 1h at room temperature with WT Spike, RBD or Nucleocapsid proteins covalently coupled to distinct carboxylated beads in a triplex assay. The liquid phase was aspirated and beads were washed with 10mM PBS/0.05% Tween-20 three times before incubation with PE-conjugated anti-human IgG-Fc antibody (Leinco/Biotrend) for 30 minutes. Beads were washed again as above and resuspended in 100μl PBS/Tween before being analysed on a Luminex analyser (Luminex/R&D Systems) using Exponent Software V31. Specific binding of antibodies to each protein was reported as the mean fluorescence intensity (MFI).

### Pseudotype neutralisation assay

Sera were heat-inactivated at 56°C for 30 min, then frozen in aliquots at -80°C. Virus neutralisation assays were performed on HEK293T cells that were transiently transfected with ACE2 using a SARS-CoV-2 Spike pseudotyped virus expressing luciferase, as previously described(42). Pseudotyped virus were prepared by transfection of HEK293T cells using the Fugene HD transfection system (Promega), as previously described(42). Pseudotyped virus was incubated with serial dilution of heat-inactivated human serum samples in duplicate for 1h at 37°C. Virus-only and cell-only controls were included. Freshly trypsinised HEK293T ACE2-expressing cells were added to each well. Following 48h incubation in a 5% CO2 environment at 37°C, luminescence was measured using the Bright-Glo Luciferase assay system (Promega) and neutralization calculated relative to virus-only controls. Neutralising antibody titres at 50% inhibition (ND_50_) were calculated in GraphPad Prism.

### Statistical analysis

Statistical analyses were carried out using GraphPad Prism v9.5.0 and in *R* version 4.2.3. Comparison of paired datasets was done using the Wilcoxon matched paired-signed ranked test to track the trending pattern between the paired samples and the Mann-Whitney U test was used to compare distributions between two groups. Multiple group differences were analysed using Kruskal-Wallis tests and pairwise differences across groups were examined using the Dunn’s test, with Benjamini-Hochberg corrections to account for potential false discovery from multiple comparisons. Linear regression models were optimised to assess relationships between variables where indicated. All p-values are two-tailed where ns>0.05, *≤0.05, **≤0.01, ***≤0.001, ****≤0.0001.

For visualisation, scatter, violin, and boxplots were generated using the ggplot2 package (version 3.4.2). Contour lines were superimposed upon a scatter plot using the geom_hdr function provided by the ggdensity package (version 1.0.0). Correlation values, including both Pearson and Spearman coefficients, were determined via the cor.test function of the stats package. These values were then annotated on the plots using the annotate function inherent to ggplot2. The summary heatmap was created with the ComplexHeatmap package (version 2.15.4).

### Data availability statement

All data generated in this study are available as supplementary information.

## Results

### Anti SARS-CoV-2 antibody profiling in Rituximab treated vasculitis patients

We identified a cohort of 14 patients with primary systemic vasculitis treated with Rituximab (RTX cohort) who had been recruited in a prospective observational cohort study investigating SARS-CoV-2 vaccine responses among renal populations(38) and had paired samples, collected at pre-specified time points, after the first, second and third vaccine doses. The patient demographics are summarised in supplementary Table 1. The majority of patients (79%) had Rituximab within 12 months prior to first vaccine dose and 64% had additional Rituximab between vaccine doses. Serological responses to wild-type SARS-CoV-2 Spike and RBD, as assessed by an accredited Luminex immunoassay 4-6 weeks after each vaccine dose, are shown in Figure 1. Overall, 57% of the cohort showed a positive anti-Spike response (median [range] MFI: 3,958 [59-22,509]) and 14% a positive anti-RBD response (median [range] MFI: 115 [46-2,437]) after the first vaccine dose. Seroconversion and Luminex MFI values increased significantly after the second vaccine dose, both against Spike (100% positive; median [range] MFI: 27,713 [9,047-32,716]; p<0.0001 Mann-Whitney test) and RBD (71% positive; median [range] MFI: 5,301 [33-31,509]; p=0.0384 Mann-Whitney test), and remained stable after the third vaccine dose (100% anti-Spike positive; median [range] MFI: 27,356 [2,398-32,662]; p=0.5007 and 64% anti-RBD positive; median [range] MFI: 19,741 [127-32,331]; p=0.2915 between second and third vaccine doses; Mann-Whitney test). Notably, four patients showed consistently negative anti-RBD responses, and 42% of sera with significant anti-Spike antibody reactivity (n=33) were negative against SARS-CoV-2 RBD (n=14). Patients who received additional Rituximab between second and third vaccine doses (n=6) did not have significantly different antibody responses compared to those who did not (median [range] MFI after third vaccine: anti-Spike; 32,552 [9,047-32,716] versus 27,356 [13984-32,662], p=0.5368, anti-RBD; 19,176 [33-31,501.8] versus 26,931 [136-32,331], p=0.4286, Mann-Whitney tests).

**Figure 1.**
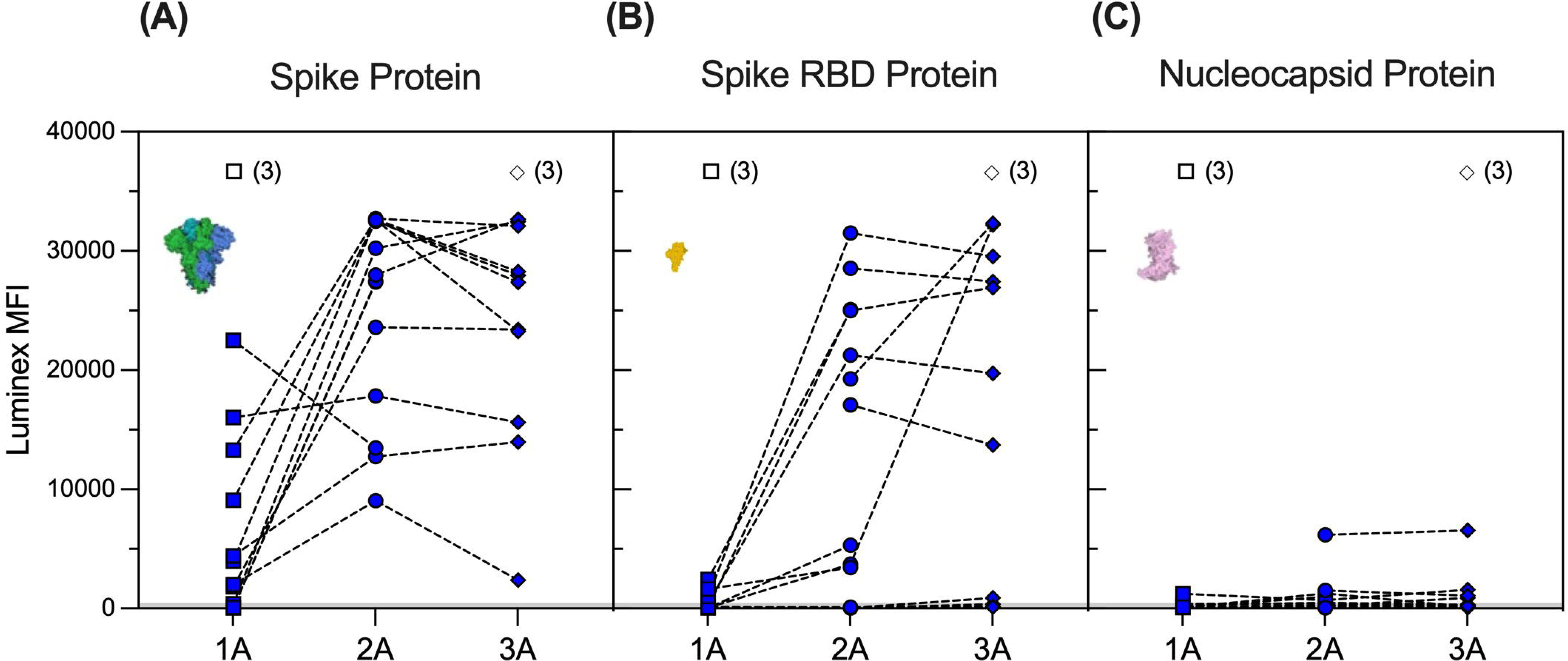
Serological responses to Wild type SARS-CoV-2 antigens in Rituximab treated patients after repeated vaccination. SARS-CoV-2 antibody profiles as determined using Luminex. The y-axis depicts mean fluorescence intensity (MFI) levels. Sera were assessed at approximately one month after 1^st^ (1A, squares), 2^nd^ (2A, circles), and 3^rd^ (3A, diamonds) vaccine dose against Spike (A), RBD (B) and Nucleocapsid (C) antigens. Symbols represent the MFI values for one sample, whereas dotted lines connect samples from the same patient taken at different timepoints. Unfilled symbols represent samples (number) where Luminex MFI data was unavailable.

Neutralising antibodies against WT were detectable in 50% of patients both after the second and third vaccine doses with, overall, similar 50% neutralising dose titres (ND_50_) between the two time points (median [range] ND_50_ of 207 [20-2,254] versus 240 [20-7,468], p=0.5890 Mann-Whitney test). Nevertheless, individual patient responses after vaccination were variable with neutralising titres increasing in six patients, decreasing in three patients, whereas a further five patients did not develop neutralising antibodies after three vaccine doses (Figure 2A). To provide a more in depth analysis of serological responses in these patients, we further characterised anti-RBD antibodies by MAAP. Consistent with Luminex, MAAP showed that the affinity (K_D_) and concentration ([Ab]) of antibodies against WT RBD did not change significantly between the second and third vaccine doses (Figure 2B-C). All neutralising sera were quantifiable by MAAP whereas non-neutralising sera could not be quantified either due to undetectable/low anti-RBD antibody levels (n=10, RBD MFI 33-3,440) or the inability to effectively constrain MAAP parameters (n=3, RBD MFI 5,301-19,741; Supplementary Figure 1 and Supplementary Table 2; one non-neutralising serum had a missing MFI and showed minimal binding on MAAP). Overall, the affinity of quantifiable anti-WT RBD antibodies ranged widely from 2.3 nM to 44.8 nM and in patients with increasing neutralisation titres after the third vaccine dose, this was driven predominantly by an increase in antibody concentration rather than improvement in affinity.

**Figure 2.**
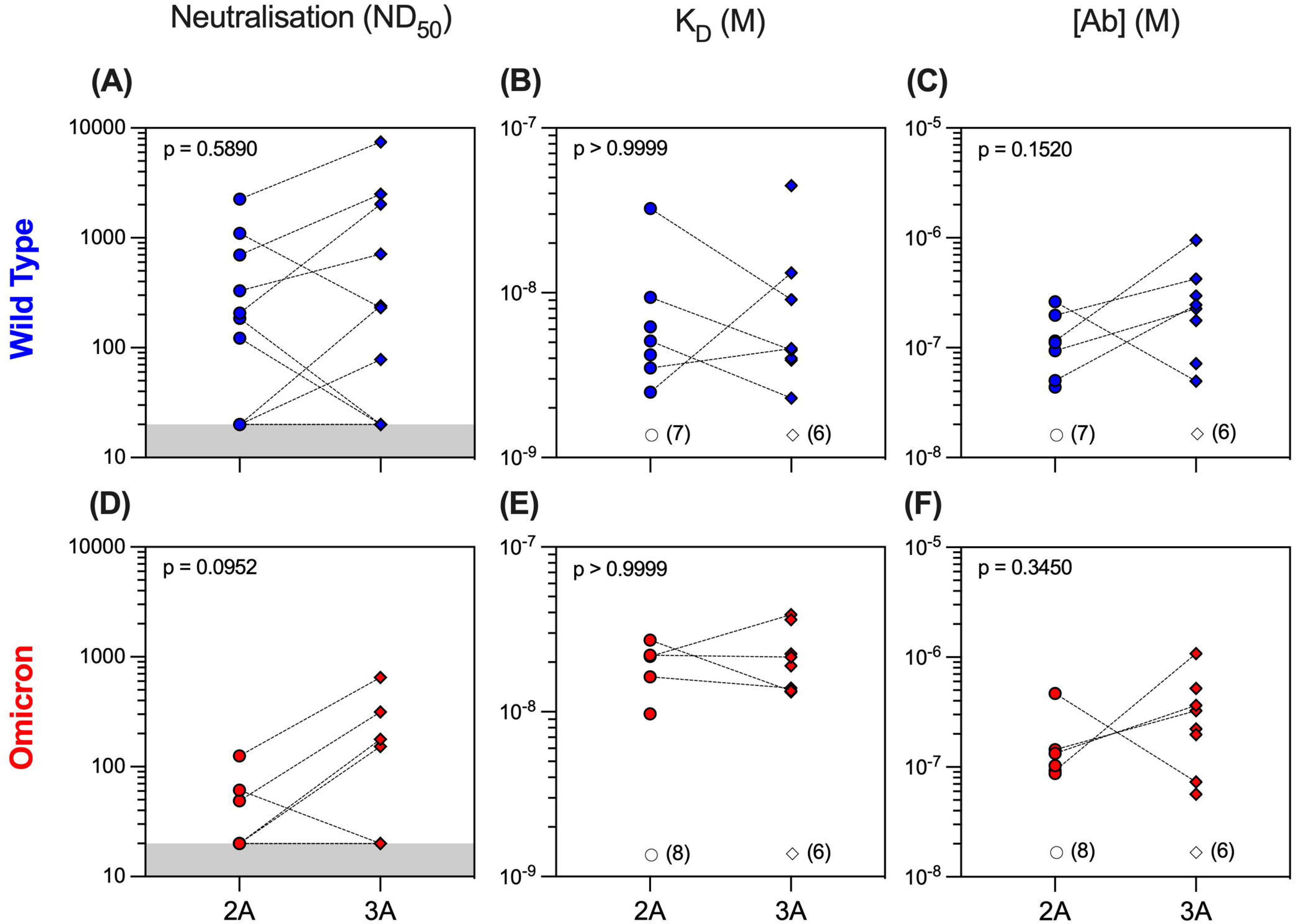
Serum neutralisation capacity and microfluidic antibody affinity profiling against Wild type and Omicron variants in Rituximab treated patient cohort. Sera from Rituximab treated vasculitis patients were assessed at one month after 2^nd^ (2A, circles, n=14) and 3^rd^ (3A, diamonds, n=14) vaccine doses for their neutralising capacity (ND_50_: 50% neutralising dose titre) against both Wild type **(A)** and Omicron **(D)** variants. All sera were quantified using microfluidic antibody affinity profiling (MAAP) to measure the affinity (K_D_, M) and concentration ([Ab], M) of antibody binding sites that specifically bind either the Wild Type **(B-C)** or Omicron **(E-F)** SARS-CoV-2 Spike RBD domains. There were no significant differences between time points for any of the comparisons using either paired (Wilcoxon paired signed ranked test) or unpaired (Mann-Whitney) statistical assessments (p-values from Man-Whitney tests are shown). Dotted lines connect samples taken from the same patient at different timepoints. Grey regions represent the lower limit of the neutralisation assay. Unfilled symbols represent the samples (number) where MAAP data was unobtainable due to non-binding or unquantifiable binding.

Analysis of serum antibody reactivity against the Omicron variant showed that only 3/14 (21%) and 4/14 (29%) patients developed neutralising antibodies after the second and third vaccine doses, respectively. Similar to WT, individual patient responses varied although more patients had measurable Omicron anti-RBD antibodies after the third vaccine dose (seven patients had an improved and three patients a worse response to Omicron RBD), as determined by MAAP (Figure 2 D-F and Supplementary Table 2). Overall, the affinity of anti-Omicron RBD antibodies varied from 9.7-38.8 nM and their concentration from 56.5-1,080 nM with no significant difference in either parameter between the two time points. Importantly, there was decreased neutralisation capacity against Omicron compared to WT both after the second and third vaccine doses (Figure 3A-C). MAAP analysis showed that this difference was driven by significantly weaker antibody affinity to the Omicron versus WT RBD (median [range] K_D_: 21.6 [9.7-38.8] nM vs 4.6 [2.3-44.8] nM respectively, p=0.0004 Wilcoxon paired ranked test) rather than by differences in antibody concentration (median [range] [Ab]: 171 [56.5-1,080] nM vs 177 [43.7-952] nM respectively, p=0.2412, Wilcoxon paired ranked test), and this was consistent at both time points (Supplementary Figure 2).

**Figure 3.**
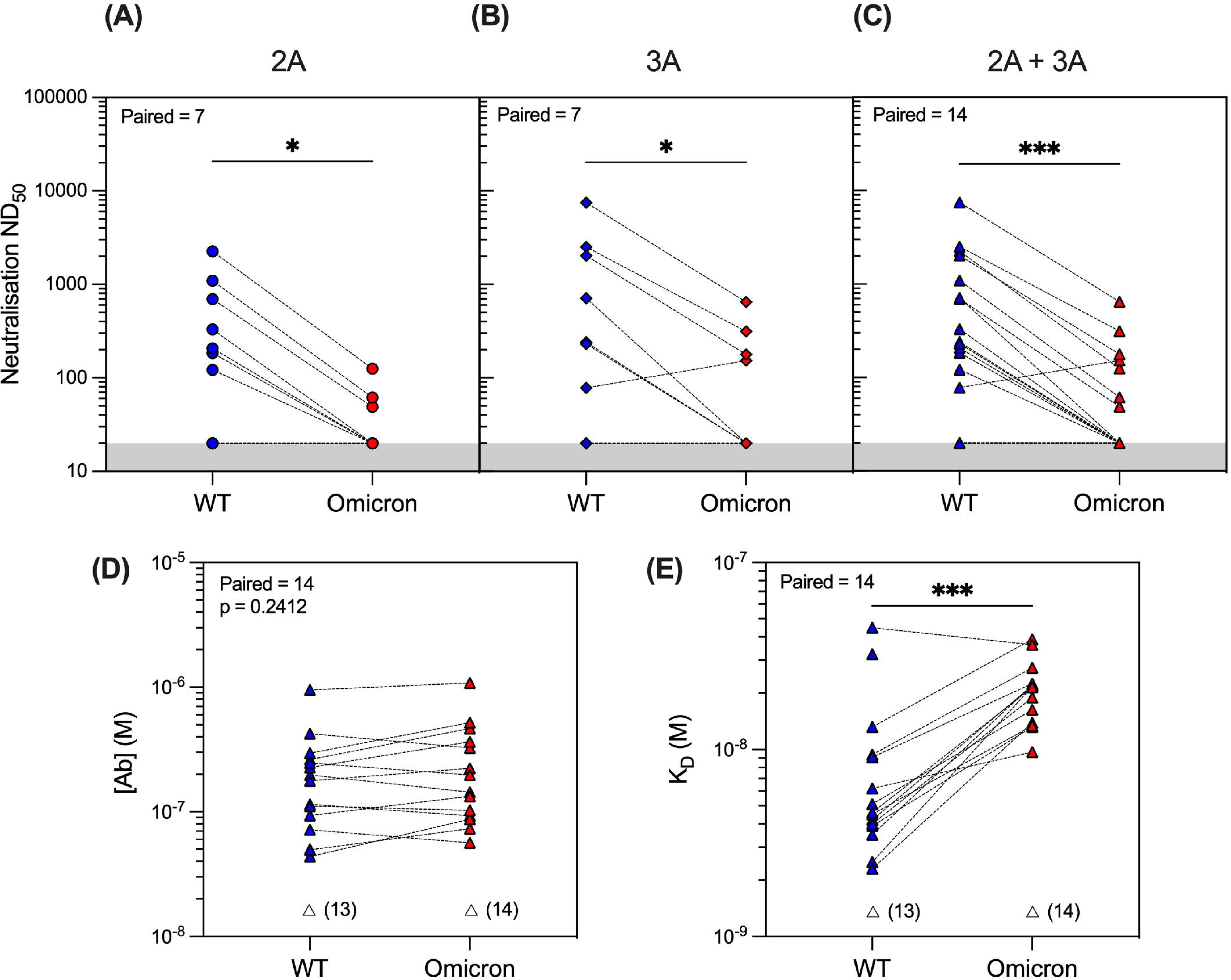
SARS-CoV-2 antibody profiling in Rituximab treated patients showed reduced neutralising capacity against the Omicron compared to Wild type (WT) variants driven by weaker antibody affinity to the Omicron Spike RBD. Neutralising capacity (ND_50_) of serum from Rituximab treated vasculitis patients against the Omicron variant (red symbols) compared to the WT variant (blue symbols) at one month after the 2^nd^ vaccine dose (**A**; 2A, circles, p=0.0156), 3^rd^ vaccine dose (**B**; 3A, diamonds, p=0.0312), or when data from both timepoints were combined (**C**; 2A+3A, triangles, p=0.0002). Microfluidic antibody affinity profiling to measure antibody affinity (K_D_, M) and binding site concentration ([Ab], M) demonstrated no difference in the concentration of antibodies recognising the WT and Omicron RBD variants (**D**, p=0.2412), but weaker antibody affinity against the Omicron strain (**E**, p=0.0004). Dotted lines connect identical samples assessed against different variants. Unfilled symbols represent the samples (number) where MAAP data was unobtainable due to insufficient or unquantifiable binding. P-values presented are two-tailed from the Wilcoxon paired signed-ranks test.

Overall, 14 (50%) samples had no neutralisation against either variant, whereas five patients did not develop neutralising antibodies at either timepoint. Taken together, the above results demonstrate that Rituximab treated patients developed widely variable antibody responses to SARS-CoV-2 after repeated vaccination which, when quantifiable by MAAP, varied by 20-fold in affinity. Overall, neutralising antibody responses were often absent even after three vaccine doses; when present, neutralisation capacity was significantly weaker against Omicron versus WT SARS-CoV-2 strains and this difference was driven by the weaker affinity of vaccine induced antibodies against the Omicron strain.

### Anti-SARS-CoV-2 antibody profiling in healthy individuals versus Rituximab treated patients

We next hypothesised that the quality of the antibody response would vary significantly in healthy individuals compared to Rituximab treated patients following antigenic exposure to SARS-CoV-2. To investigate this, we profiled antibody responses in health care workers (HCW) recruited at Cambridge University Hospital who, similar to the vasculitis cohort, had three exposures to SARS-CoV-2 antigen, consisting of an asymptomatic infection to SARS-CoV-2 prior to the emergence of the Omicron variant and two subsequent vaccine doses (see methods and supplementary Table 1). Antibody profiling was performed at 4 weeks after the third exposure (second vaccine dose). Luminex analyses showed significantly higher antibody reactivity to WT Spike and RBD in HCW versus Rituximab treated patients (Supplementary Figure 3, Supplementary Table 4). Similar to Rituximab treated patients, neutralisation titres in HCW were significantly higher against the WT versus the Omicron variant. Further profiling by MAAP suggested this difference was driven by the fact only a fraction of high affinity serum antibodies recognised the Omicron RBD (Supplementary Figure 3). Compared to Rituximab treated patients (Figure 4), MAAP quantifiable antibodies to WT SARS-CoV-2 in the HCW cohort were of higher concentration (median [range] [Ab] of 236.5 [49.5-952] nM vs 646 [233-2,110] nM, respectively, p=0.0018 Mann-Whitney test) but of similar affinity (median [range] K_D_ of 4.6 [2.3-44.8] nM vs 4.7 [1.2-19.0] nM, respectively, p=0.5825 Mann-Whitney test) underpinning the higher neutralisation titres in the HCW cohort (median [range] ND_50_ of 49 [20-7,468]) versus 2,522 [864-7,297], p=0.0004 Mann-Whitney test; Figure 4). Neutralisation titres against the Omicron strain were also higher in HCW versus Rituximab treated patients (median [range] ND_50_ of 221.5 [24.63-1,225] versus 20 [20-648.7], respectively, p=0.003 Mann-Whitney test; Figure 4) but this was driven by higher affinity anti-Omicron RBD antibodies in the HCW cohort (median [range] K_D_: 1.05 [0.45-1.84] nM vs 20.25 [13.2-38.8] nM, p=0.0002 Mann-Whitney test). Taken together, our results confirm the previously reported impaired antibody responses to SARS-CoV-2 variants of concern in Rituximab treated, immunosuppressed patients compared to healthy individuals and, through MAAP analysis, highlight the significance of the fundamental parameters of the antibody response, namely antibody affinity and concentration, to anti-viral immunity and virus escape.

**Figure 4.**
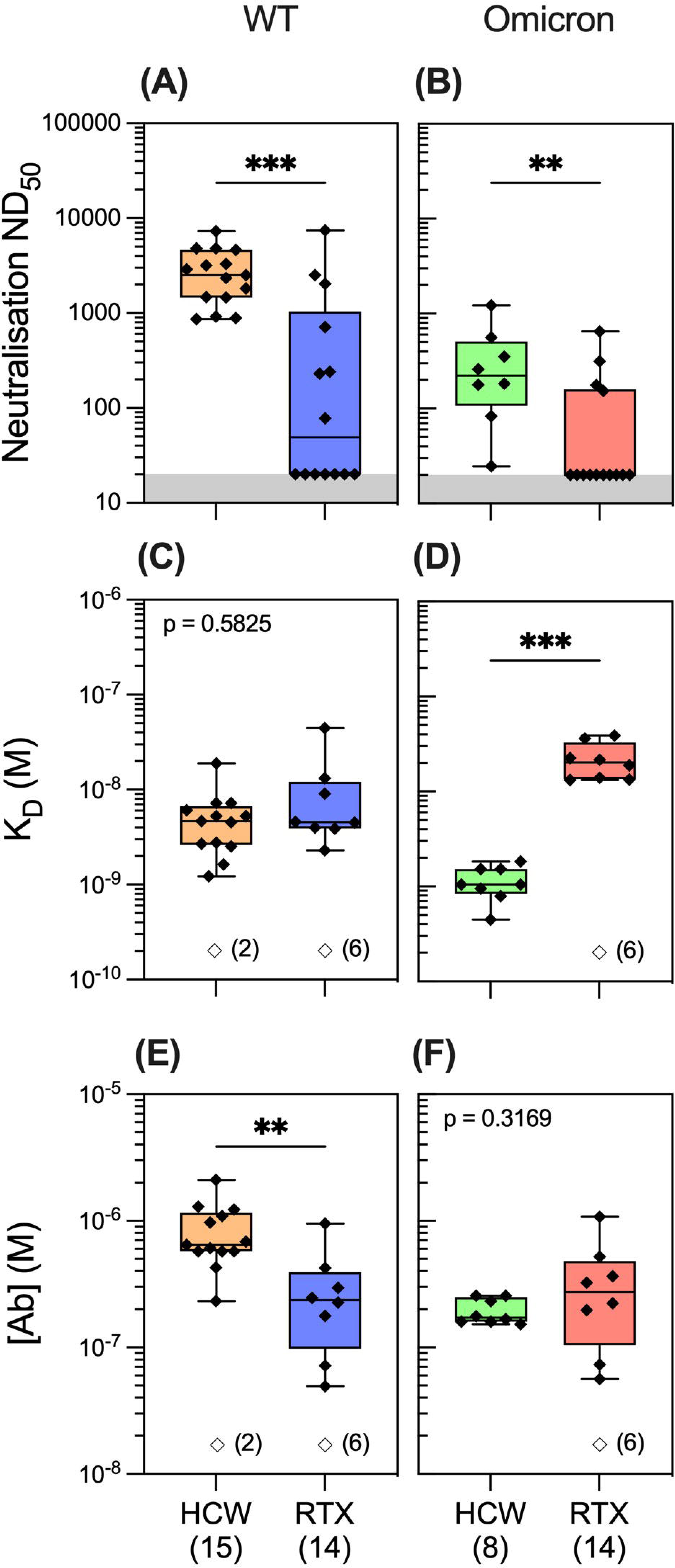
Anti-SARS-CoV-2 antibody profiling in healthy individuals versus Rituximab treated patients. Serum neutralising titres (ND_50_) in healthcare workers (HCW) versus Rituximab (RTX) treated vasculitis patients one month after the third exposure to SARS-CoV-2 antigen (3A) assessed against Wild type (**A**; p=0.0004) and Omicron strains (**B**; p=0.003). Comparison of quantifiable antibodies to Wild type SARS-CoV-2 using microfluidic affinity profiling (MAAP) showed similar affinity (**C;** p=0.5825), but lower abundance in Rituximab treated patients (**E**; p=0.0018). In contrast, the abundance of antibodies against the RBD Omicron variant was comparable between the cohorts (**F**; p=0.3169), however these antibodies were of higher affinity in the HCW cohort (**D**; p=0.0002). In **A-B**, the grey region represents the lower limit of detection for the neutralisation assay. Box blots represent the median, range and interquartile range for each dataset. Unfilled symbols in **C-F** represent samples (number) where MAAP data was unobtainable due to insufficient or unquantifiable binding. Statistical analysis was carried out for each plot via Mann Whitney Test and presented p-values are two-tailed.

### Evolution of antibody response to SARS-CoV-2 after infection and vaccination

In the microfluidic-based antibody profiling of the Rituximab patient cohort described above, we could not detect evidence of affinity maturation in peripheral blood samples collected after the second compared to third vaccine doses, despite a median of 185 days (range 141-224 days) between the two time points. To investigate whether this observation was specific to the Rituximab treated patients, we next examined the quality of the humoral response after WT SARS-CoV-2 infection in patients during the first wave of the COVID-19 pandemic using samples collected at one and three months post infection (see methods). The demographic characteristics of this cohort (n=30) are shown in supplementary Table 1. As shown in Figure 5, we observed wide variation in affinity (K_D_ range 2.07-34.0 nM) and concentration (range 7.1-1,120 nM) of the antibody responses to WT RBD at both time points. As expected, there was a significant decrease in antibody concentration over time (median [range]: 55.6 [7.1-269] nM at 3 months vs 169 [13-1,120] nM at 1 month post-infection, p=0.0034, Wilcoxon paired ranked test) and an associated decrease in neutralisation capacity (median [range] ND_50_ of 131 [20-7,983] versus 280 [20-14,580], respectively, p=0.0296, Wilcoxon paired ranked test). Nevertheless, we were able to detect evidence of affinity maturation over the same time period, as demonstrated by a decrease in anti-RBD dissociation constant in the majority of patients (median [range] K_D_: 6.6 [2.1-14.6] nM at 3 months vs 9.4 [2.1-34] nM at 1 month post-infection, p=0.0244, Wilcoxon paired ranked test). Luminex analysis of antibody reactivity to WT Spike and RBD showed significantly higher responses at one month after infection in the convalescent cohort compared to one month after the first vaccine dose in the Rituximab cohort; these antibodies in Rituximab treated patients reached similar levels to those in the convalescent cohort after the second and third vaccine doses (supplementary Figure 4D-E). Comparison of the convalescent cohort at one month post-infection with the Rituximab treated patient cohort at equivalent time points after the second and third vaccine dose showed similar neutralisation titres against WT SARS-CoV-2 despite exposure to only a single sensitisation event (Figure 6). Corroborating these findings, quantification of WT anti-RBD responses by MAAP showed similar antibody affinity and concentration after the second and third vaccine dose in Rituximab treated patients compared to those in post-infection convalescent patients (Figure 6B-C).

**Figure 5.**
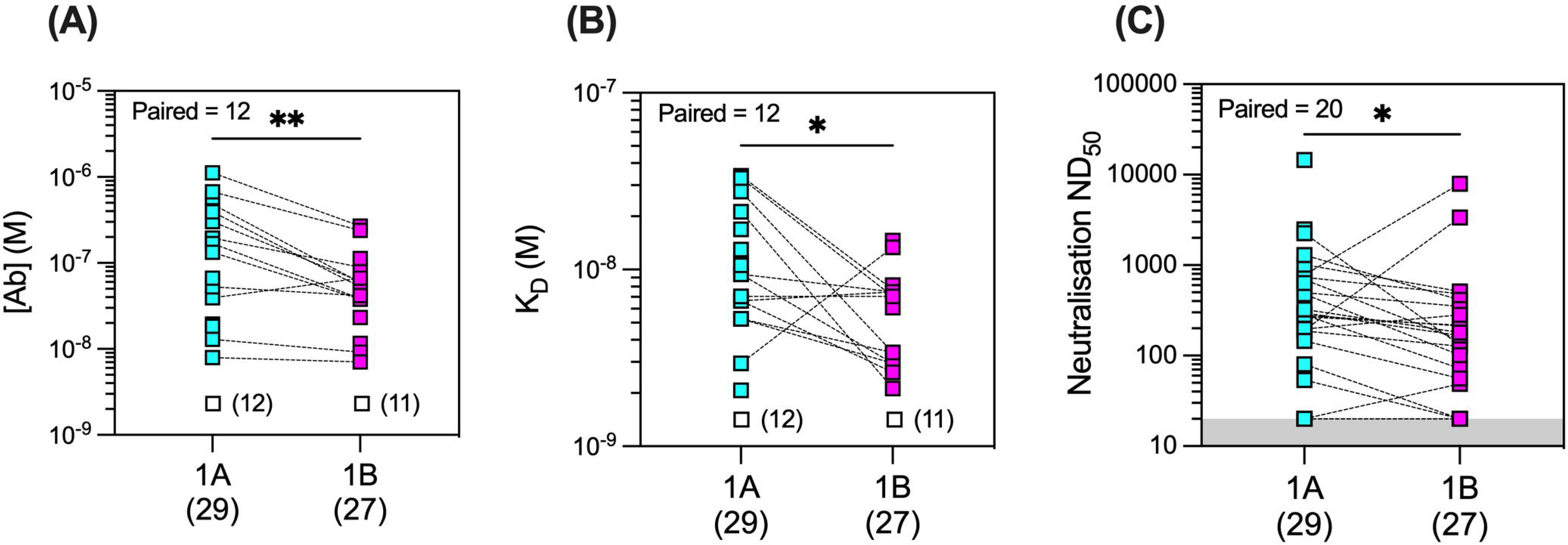
Microfluidic antibody affinity profiling (MAAP) of the evolution of the antibody response after infection with Wild type SARS-CoV-2. Anti-SARS-CoV-2 Wild type Spike RBD antibody affinity (K_D_, M) and binding site concentration ([Ab], M) in sera from convalescent patients at one-(1A) and three-months (1B) post-infection were assessed using MAAP. Panel **A** shows that the concentration of antibodies decreased over time (**A**; p=0.0034), but their affinity against RBD increased (**B**; p=0.0244), resulting in an overall decrease in serum neutralisation capacity (**C**; p=0.0296). Dotted lines connect samples taken from the same patient at the two timepoints. Unfilled symbols in **A-B** represent samples (number) where MAAP data was unobtainable due to insufficient or unquantifiable binding. Presented p-values are two-tailed from the Wilcoxon paired signed rank test.

**Figure 6.**
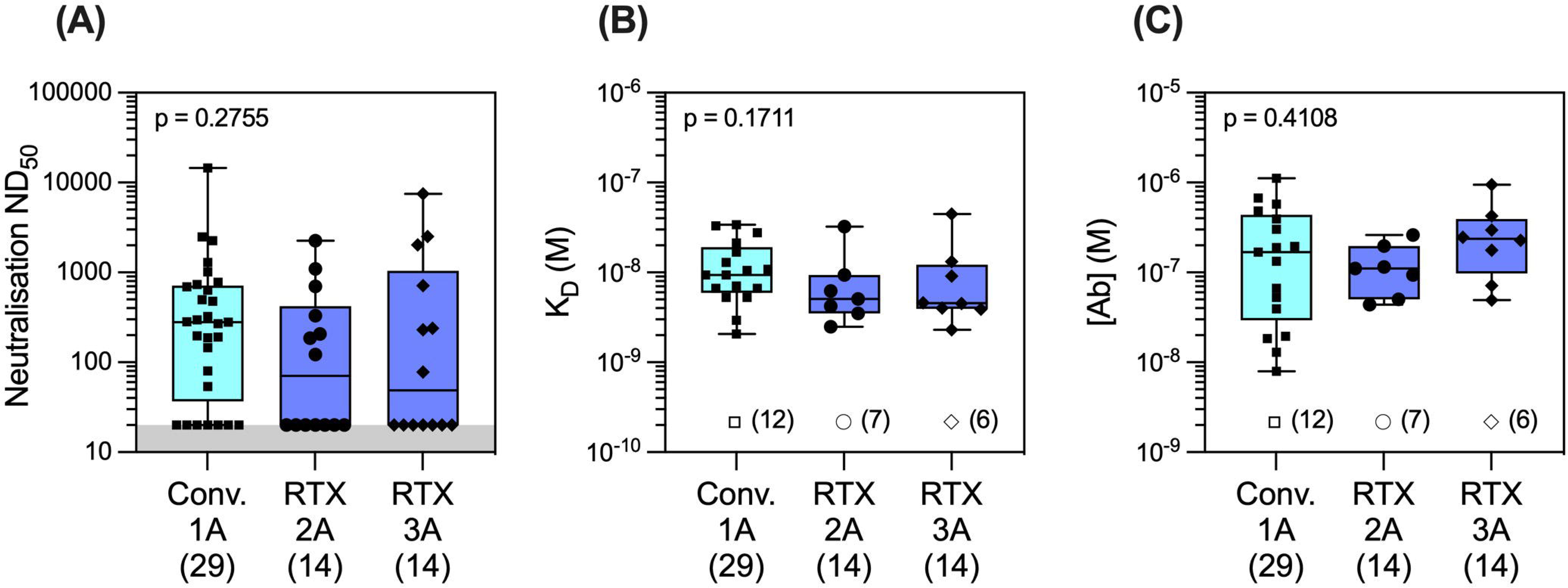
Antibody profiling against Wild type SARS-CoV-2 in Convalescent versus Rituximab treated patients. Comparison of serum neutralising titres (ND_50_) in convalescent patients one month after Wild type SARS-CoV-2 infection (Conv. 1A) versus in Rituximab treated patients at one month following the second (RTX 2A) and third vaccine doses (RTX 3A) showed numerically higher, but not statistically different titres in the two cohorts **(A;** Conv. 1A vs RTX 2A: p=0.2832 and Conv. 1A vs RTX 3A: p=0.5515). Comparison of quantifiable antibodies to Wild type SARS-CoV-2 using microfluidic affinity profiling (MAAP) showed similar antibody affinity (**B**; Conv. 1A vs RTX 2A: p=0.2236 and Conv. 1A vs RTX 3A: p=0.3002) and antibody concentration (**C**; Conv. 1A vs RTX 2A: p>0.9999 and Conv. 1A vs RTX 3A: p=0.6399). Box blots represent the median, range and interquartile range for each dataset. Grey shading in A represents the lower assay limit. Unfilled symbols in B-C represent samples (number) where MAAP data was unobtainable due to insufficient or unquantifiable binding. Statistical analysis was carried out using Kruskal-Wallis ANOVA test (presented) and Dunn’s multiple comparisons and all p-values are two-tailed.

### Multidimensional assessment of antibody fingerprints (concentration and affinity) with clinical parameters at the individual patient level

To provide a global perspective of antibody affinity/concentration profiles against WT and Omicron variants, we created integrated 2D-density contour plots incorporating MAAP data obtained from the above three patient groups (Rituximab treated vasculitis patients ∼1 month after the third vaccine dose vs health care workers ∼1 month after the third sensitisation event vs COVID-19 convalescent patients ∼1 month post-infection). As shown in Figure 7, HCW clustered separately (high affinity and high [Ab] profiles), whereas the density representations for Rituximab treated and convalescent patients were wider and largely overlapping. As discussed above, a left shift of antibody profiles to the Omicron variant was apparent in Rituximab patients with significantly lower affinity compared to anti-Omicron profiles in the HCW cohort. The relationship of antibody fingerprints with clinically relevant parameters showed that the K_D_ x [Ab] product was similar between female and male patients, whereas a weak negative correlation was noted with increasing age for anti-WT antibody profiles (Spearman correlation coefficient ρ=-0.67, p=0.0012; Figure 7B-C). For convalescent patients, the K_D_ x [Ab] product tended to be higher in patients with more severe disease (Figure 7D).

**Figure 7.**
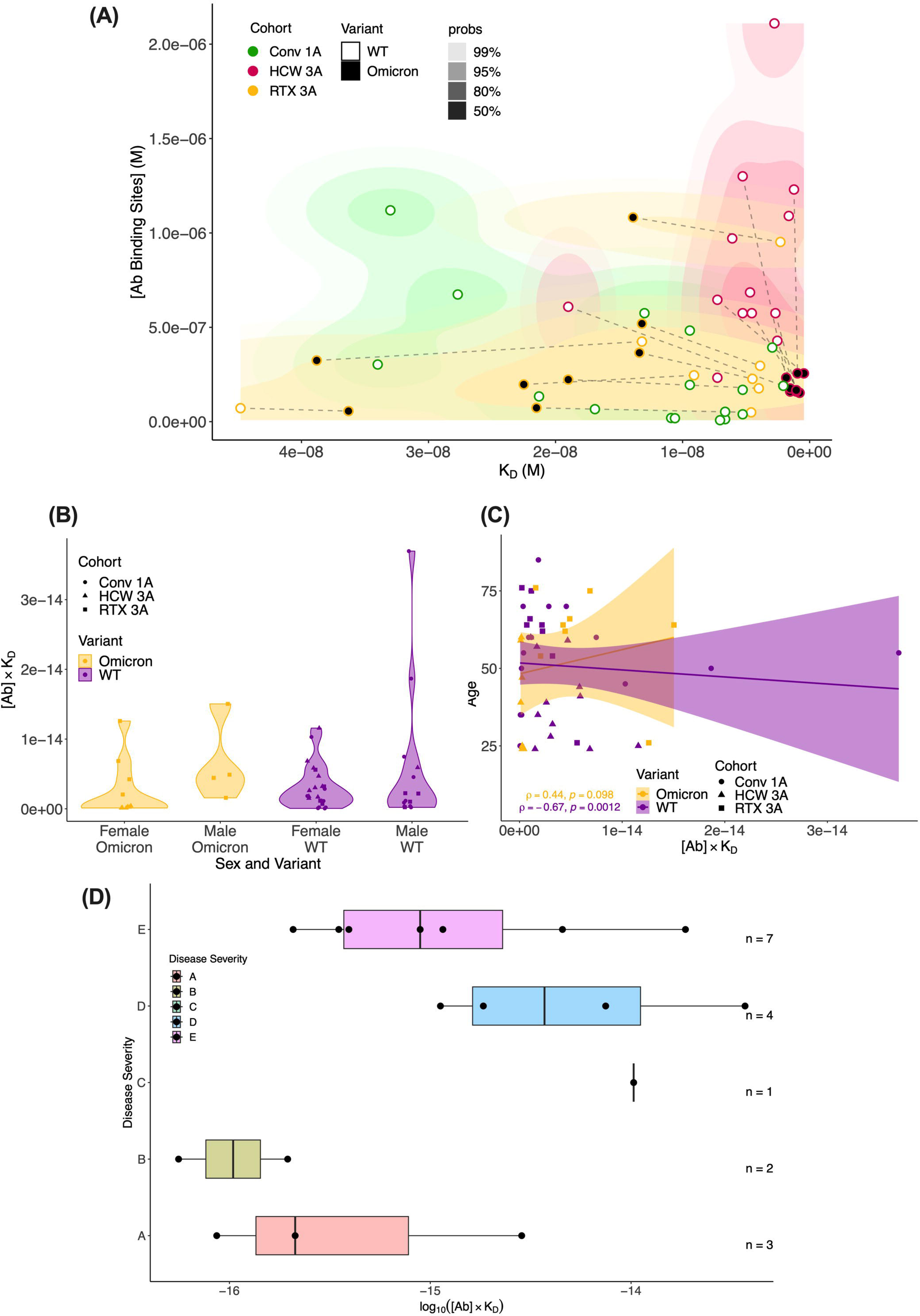
Global analysis of antibody fingerprints to SARS-CoV-2 Wild type and Omicron variants across all patient cohorts and correlation with clinically relevant parameters. **Panel A. Antibody affinity and concentration across different patient cohorts and variants.** The contour map summarises the equilibrium dissociation constant (K_D_) vs the concentration of antibody binding sites ([AB]) across three cohorts: Conv. 1A (convalescent COVID-19 1-month post-infection, green), RTX 3A (Rituximab cohort 1-month after 3rd vaccine dose, yellow), and HCW 3A (health care worker cohort 1-month after infection and 2 vaccinations, red); the colour gradient is proportional to the expected probability. The filling colour of points indicates the variant. Wild type (WT) and Omicron points from the same sample are linked by dashed lines. The x-axis depicts increased affinity i.e. lower K_D_ values towards the right; higher concentrations of antibody binding sites are displayed, in increasing order, on the y-axis. **Panel B. Comparison of antibody affinity and concentration across sex and SARS-CoV-2 variants.** The plots summarise the product of equilibrium dissociation constant (K_D_) and concentration of antibody binding sites ([AB]) in relation to sex and SARS-CoV-2 variants (WT and Omicron) across three cohorts: Conv. 1A (convalescent COVID-19 1-month post-infection, circles), RTX 3A (Rituximab cohort 1-month after 3rd vaccine, squares), and HCW 3A (health care worker cohort 1-month after infection and 2 vaccinations, triangles). On the x-axis, the multiplicative classes on sex (male or female) and variant are represented. The y-axis represents the product of K_D_ and [AB] (no units). No significant differences in the product of antibody affinity and concentration were noted between female and male individuals for both Omicron (p=0.0617) and WT (p=0.0539); p-values are from Mann-Whitney U test. **Panel C. Comparison of antibody affinity and concentration according to patient age and SARS-CoV-2 variant.** This scatter plot with a linear regression line depicts the interaction between age (y-axis) and the product of the equilibrium dissociation constant (K_D_) and concentration of antibody binding sites ([AB]) (x-axis), across the variants (WT and Omicron) and cohorts: Conv. 1A (convalescent COVID-19 1-month post-infection, circles), RTX 3A (Rituximab cohort 1-month after 3rd vaccine, squares), and HCW 3A (health care worker cohort 1-month after infection and 2 vaccinations, triangles). A linear model fit (smooth line) and the Spearman rank correlation coefficient is depicted, indicating the strength and direction of the correlation between age and [AB]xK_D_. **Panel D. Relationship between COVID-19 disease severity and the product of antibody affinity and binding site concentration (log(AB*K_D_) at 1-month post-infection in the convalescent cohort.** Convalescent patients after more severe disease showed a trend towards higher antibody responses ([AB]xK_D_) to WT SARS-CoV-2 one month after infection compared to asymptomatic patients and those with mild disease (Kruskal-Wallis rank sum test: p=0.08; pairwise comparisons with Benjamini-Hochberg multiple testing correction: p>0.05 for all comparisons). Disease severity was classified as follows: (A) asymptomatic; (B) mild symptoms not requiring hospitalisation; (C) patients who presented to hospital but never required oxygen supplementation; (D) patients who were admitted to hospital and whose maximal respiratory support was supplemental oxygen; and (E) patients who at some point required assisted ventilation.

We next incorporated serological parameters (MAAP affinity/concentration, Luminex MFI, ND_50_) with relevant immunological and demographic variables (SARS-CoV-2 variant, vaccine dose, patient cohort, age, sex and ethnic background) to provide a comprehensive multidimensional assessment at the single-individual level (Figure 8A), which highlighted the strength of the anti-SARS-CoV-2 response (higher ND_50_, [Ab], Luminex anti-Spike and anti-RBD MFI) in healthy individuals, compared to Rituximab treated and convalescent patients. We also related antibody WT SARS-CoV-2 serological read-outs obtained from neutralisation, MAAP and Luminex assays, which showed a strong correlation between ND_50_ and antibody concentration or ND_50_ and Luminex RBD MFI (Spearman correlation coefficient ρ=0.74, p=8.1×10^-12^ and ρ=0.81, p=5.5×10^-24^, respectively; Figure 8B-C) but no relationship between ND_50_ or Luminex RBD MFI with antibody affinity (supplementary Figure 5). These observations suggest that neutralisation and solid-phase Luminex assays primarily reflect antibody concentrations (supplementary Figure 5C) rather than affinities for MAAP quantifiable samples analysed here.

**Figure 8.**
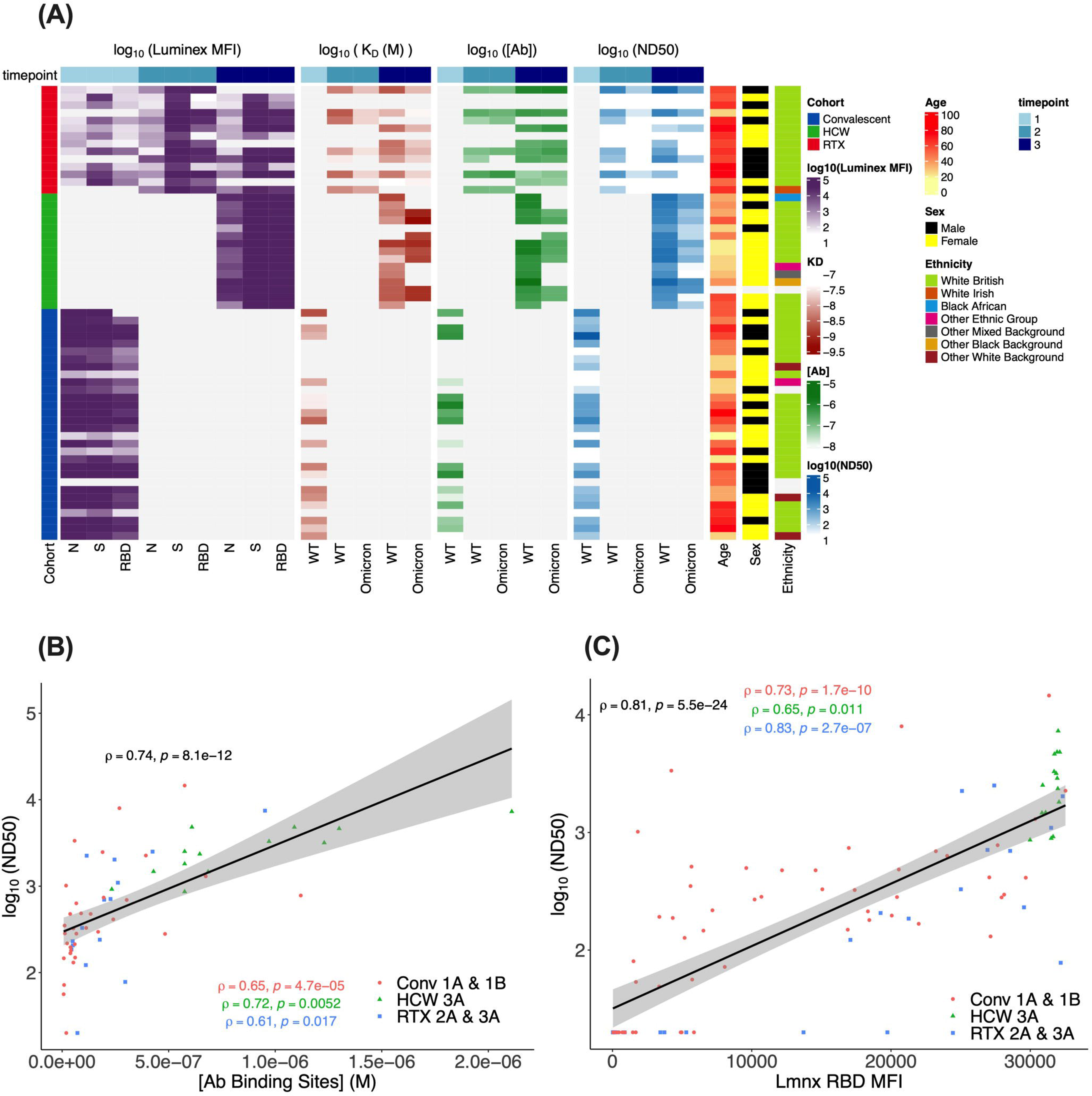
Multidimensional assessment of SARS-CoV-2 antibody profiles at the individual patient level. **Panel A. Summary heatmap depicting key immunological and demographic metrics across different cohorts and timepoints.** Each row corresponds to a single participant in the study, grouped per cohort affiliation (Left hand side colour bar). Columns correspond to several measurements, structured into four main categories: Luminex MFI, K_D_, [Ab], and ND_50_. Each of these categories contains measurements at three distinct time points which are denoted by the top colour bar. Luminex MFI has a breakdown into N, S, and RBD. For K_D_, [Ab], and ND_50_, measurements are differentiated by the variant of the virus (WT or Omicron). Colour intensity within the heatmap corresponds to the log-transformed value of each measurement, with darker hues indicating higher values (except in KD as lower values correspond to higher antibody affinity). Age, sex, and ethnicity of participants are also visualised in the right-hand side columns. **Panel B. Relationship between serum Wild type SARS-CoV-2 neutralisation titre (logND_50_) and antibody binding site concentration ([Ab}, M) across timepoints and patient cohorts.** Data points from each patient cohort are represented by different shapes. A linear model fit (smooth line) is shown and the Spearman correlation coefficient is depicted, indicating the strength and direction of the correlation between [AB] and ND_50_. The Spearman correlation coefficients for data in each patient cohort are also shown. **Panel C. Relationship between serum Wild type SARS-CoV-2 neutralisation titre (logND_50_) and Luminex anti-RBD MFI across patient cohorts.** Data points from each patient cohort are represented by different colours. A linear model fit (smooth line) is shown and the Spearman correlation coefficient is depicted, indicating the strength and direction of the correlation between ND_50_ and Luminex RBD MFI. The Spearman correlation coefficients for data in each patient cohort are also shown.

## Discussion

In this study, primary systemic vasculitis patients treated with Rituximab had significantly impaired humoral immune responses compared to healthy individuals after three exposures to SARS-CoV-2 antigens, demonstrated by lower anti-Spike and -RBD antibody levels and lower neutralisation titres against both wildtype and Omicron variants. Despite an incremental increase in antibody levels after the second vaccine dose, no further enhancement of the response was noted after a third dose. Correspondingly, after a third vaccine dose, approximately half and two thirds of Rituximab patients had no detectable neutralising antibodies against WT and Omicron, respectively. Relative to unvaccinated convalescent individuals, Rituximab treated patients had similar anti-Spike and anti-RBD antibody levels and similar (albeit numerically lower) neutralisation titres to WT SARS-CoV-2, despite two more exposures to SARS-CoV-2 antigen. Antibodies produced by Rituximab treated patients varied by 20-fold of magnitude in affinity, and the difference in neutralisation capacity between Omicron and WT SARS-CoV-2 was driven by the weaker affinity of vaccine induced antibodies against the Omicron variant. By contrast, healthy individuals with hybrid immunity in the form of previous infection and two vaccinations produced a broader, high concentration antibody response as profiled using MAAP, a subset of which recognised Omicron RBD with higher affinity than antibodies in Rituximab treated patients, likely underpinning the more effective neutralisation capacity against Omicron in this group compared to Rituximab treated patients. Finally, we demonstrate that MAAP can assess the evolution of humoral immune responses by direct measurement of polyclonal antibody affinity in serum, without the need for antigen-specific B cell isolation and antibody sequencing or measurement of antibody dissociation kinetics in solid-phase assays that can be perturbed by avidity-driven interactions(43–45). Accordingly, we detected evidence of affinity maturation within three months from primary infection in convalescent patient sera despite a relative decrease in anti-RBD antibody concentration. This was not evident in Rituximab treated patients even after a third vaccine dose.

Our findings of impaired humoral immune responses after a third SARS-CoV-2 vaccine dose in our primary systemic vasculitis patient cohort receiving anti-CD20 therapies are in agreement with the recently published literature, although we are unaware of any other groups who have quantified these using a combination of neutralisation assays and MAAP. A recently published study of 21 patients with antineutrophil cytoplasmic antibody vasculitis (AAV) showed that none of their 8 Rituximab-treated patients had detectable neutralising antibodies to B.1.617.2 (Delta) following a third dose of BNT162b(35). By contrast, in a small cohort of 15 AAV patients on Rituximab who received a third vaccine dose, 7/15 (46.7%) developed measurable IgG antibodies to the S1 subunit of the SARS-CoV-2 Spike protein, although the neutralisation capacity of these antibodies was not assessed(36). Humoral immune responses following a third vaccine dose among patients on anti-CD20 therapies with other autoimmune/inflammatory conditions are also impaired compared to similar patients on other immunosuppressive agents and healthy controls(14–16, 31, 46, 47). Additionally, among patients on anti-CD20 therapies who had not seroconverted following primary vaccination, a third dose led to seroconversion rates of approximately 15-60%(29, 32–34, 48, 49). Another small study demonstrated an inverse correlation between Rituximab dose and seroconversion and that antibody levels persisted after a subsequent dose of Rituximab among those who had already seroconverted(50). A fall in antibody levels following a dose of Rituximab between a second and third SARS-CoV-2 vaccine dose was recently reported, although neutralisation capacity was preserved in seroconverted patients(51). In an open-label study of a fourth dose of mRNA vaccine among Rituximab treated patients, predominantly those with rheumatoid arthritis or connective tissue diseases, a moderate improvement in seroconversion from 33% to 58% was demonstrated after the fourth dose(30).

Among the patients in our cohort with detectable anti-Spike antibody responses following SARS-CoV-2 vaccination, the impact of Rituximab on B cell populations, particularly memory B cells, may be of relevance to their impaired neutralising ability and affinity maturation compared to healthy controls. Memory B cells, in addition to enabling anamnestic antibody responses, also allow for the development of reactive humoral responses in the event of exposure to variant pathogens(52, 53). After vaccination, SARS-CoV-2 specific memory B cells have been shown to persist for several months, even after antibody titres have declined(54). Furthermore, a third dose of mRNA vaccine has been shown to increase the breadth and potency of memory B cells and their antibody responses, including against Omicron(54, 55). Following Rituximab treatment, memory B cells are often the last to reconstitute among peripheral blood B cell populations(56–59), a process that can take up to several years(57). Additionally, Rituximab treatment has also been associated with delays in the acquisition of somatic hypermutations among repopulated memory B cells(60). Recent data suggest that the presence of detectable peripheral circulating B cells is critical for seroconversion following vaccination among Rituximab treated patients(29), with SARS-CoV-2 specific memory B cell and plasmablast populations positively correlating with antibody titres and neutralisation(61). Furthermore, a preponderance of naïve and transitional B cells among Rituximab-treated patients prior to vaccination, indicative of early B cell reconstitution, was found to be associated with adequate humoral immune responses following vaccination(62). In support of this, improved serological responses after SARS-CoV-2 vaccination have been associated with increasing time since the last Rituximab dose, particularly if this interval is more than six months(63, 64). Notably in our cohort, 36% of patients received Rituximab within 6 months from the first vaccine dose and 64% received an additional Rituximab dose between first and third vaccine dose potentially accounting for the lack of improvement in the humoral response and the absence of affinity maturation between second and third vaccine doses.

A strength of our study is that we prospectively evaluated antibody responses at specific time points in a relatively understudied cohort of primary systemic vasculitis patients who are particularly vulnerable to both poorer clinical outcomes from COVID-19 and impaired humoral immune responses after vaccination. Importantly, by quantifying antibody affinity and concentration directly in serum, together with data from solid-phase Luminex and neutralisation assays, we were able to provide additional insights into the quality of the immune response against WT and Omicron variants after vaccination in Rituximab treated patients. Additionally, we provide a comparative assessment with the humoral response at equivalent time points in non-immunocompromised cohorts after infection/vaccination and after infection only. In this regard, this is the largest study to date using a novel microfluidic assay to quantify SARS-CoV-2 antibody affinity and concentration in serum samples. The limitations of our investigations reside in the relatively small number of Rituximab treated patients included in the study and the lack of a pre-Omicron control group of previously uninfected healthy individuals with three vaccinations. It would have also been informative to incorporate analysis on antigen-specific T cell and B cell mediated cellular immunity and how this may relate to the observed serological response. Finally, the limits of sensitivity of the MAAP assay should be noted. These reflect the lower limit of labelled antigen concentration that can be employed owing to intrinsic background serum fluorescence and the fact that antibody affinity and concentration cannot be constrained in samples with low concentration and/or low affinity antibodies (e.g. where [Ab] < K_D_)(25).

In conclusion, our results confirm and enrich the previously reported observations of impaired humoral immune responses to SARS-CoV-2, including variants of concern, in Rituximab treated, immunosuppressed patients compared to healthy individuals. Through the addition of MAAP analysis, we highlight the significance of the fundamental parameters of the antibody response, namely antibody affinity and concentration, to anti-viral immunity and viral escape. Consequently, should our results be replicated, we would caution against interpreting the presence of solid-phase assay detected anti-Spike antibodies following vaccination as providing evidence of immune protection against SARS-CoV-2 infection, particularly among patients on anti-CD20 therapies.

## Acknowledgements

This study was funded in part by the University of Cambridge’s Wellcome COVID-19 Rapid Response DCF (RG93172 to VK). We acknowledge funding support from the National Institute for Health Research Blood and Transplant Research Unit (NIHR BTRU, Grant number: NIHR203332) in Organ Donation and Transplantation at the University of Cambridge in collaboration with Newcastle University and in partnership with NHS Blood and Transplant (NHSBT). The views expressed are those of the authors and not necessarily those of the National Health Service, the National Institute for Health Research, the Department of Health or National Health Service Blood and Transplant. VK acknowledges funding from an NIHR Fellowship (PDF-2016-09-065) and as a Paul I. Terasaki Scholar. RS acknowledges funding from Addenbrooke’s Charitable Trust and Vasculitis UK. RK and IM are funded by the Wellcome Trust [203151/Z/16/Z]. We thank all the patients and health care workers who consented to take part in this study. We also thank the NIHR Cambridge Biomedical Research Centre for support with sample recruitment. For the purpose of open access, the authors applied a CC BY public copyright licence to all versions of the manuscript arising from this submission.

## Author contributions

AP, RS, TPJK and VK contributed to conception and design of the study. AP, SM, GM and CKX performed MAAP experiments and analysed the respective data. MXC-X, RJ, JRB and DJC collected and annotated patient samples. MH and IGG performed neutralisation studies and RD performed Luminex analyses. AP, RK and IIM performed statistical analyses. MXC-X, AP and VK wrote the manuscript. All authors read and revised the article and approved the submitted version.

*Declaration of interests*

