## Supplementary for "Microfluidic antibody profiling after repeated SARS-CoV-2 vaccination links antibody affinity and concentration to impaired immunity and variant escape in patients on anti-CD-20 therapy"

[1]Department of Surgery, University of Cambridge, Addenbrooke's Hospital, Hills Road, Cambridge CB2 0QQ, UK. [2]Department of Medicine, University of Cambridge School of Clinical Medicine, Cambridge, UK and Cambridge University Hospitals NHS Foundation Trust, cambridge, UK. [3]Centre for Misfolding Diseases, Yusuf Hamied Department of Chemistry, University of Cambridge, Lensfield Road, Cambridge CB2 1EW, UK. [4]Department of Pathology, Division of Virology, University of Cambridge, Cambridge, UK. [5]Wellcome-MRC Cambridge Stem Cell Institute, Jeffrey Cheah Biomedical Centre, University of Cambridge, CB2 0AW, UK. [6]Department of Clinical Biochemistry and Immunology, Addenbrooke's Hospital, Cambridge CB2 0QQ, UK. [7]Cavendish Laboratory, Department of Physics, University of Cambridge, JJ Thomson Ave, Cambridge CB3 0HE, UK. [8]NIHR Blood and Transplant Research Unit in Organ Donation and Transplantation at the University of Cambridge and the NIHR Cambridge Biomedical Research Centre, Cambridge, UK

### 1 Supplementary Figures

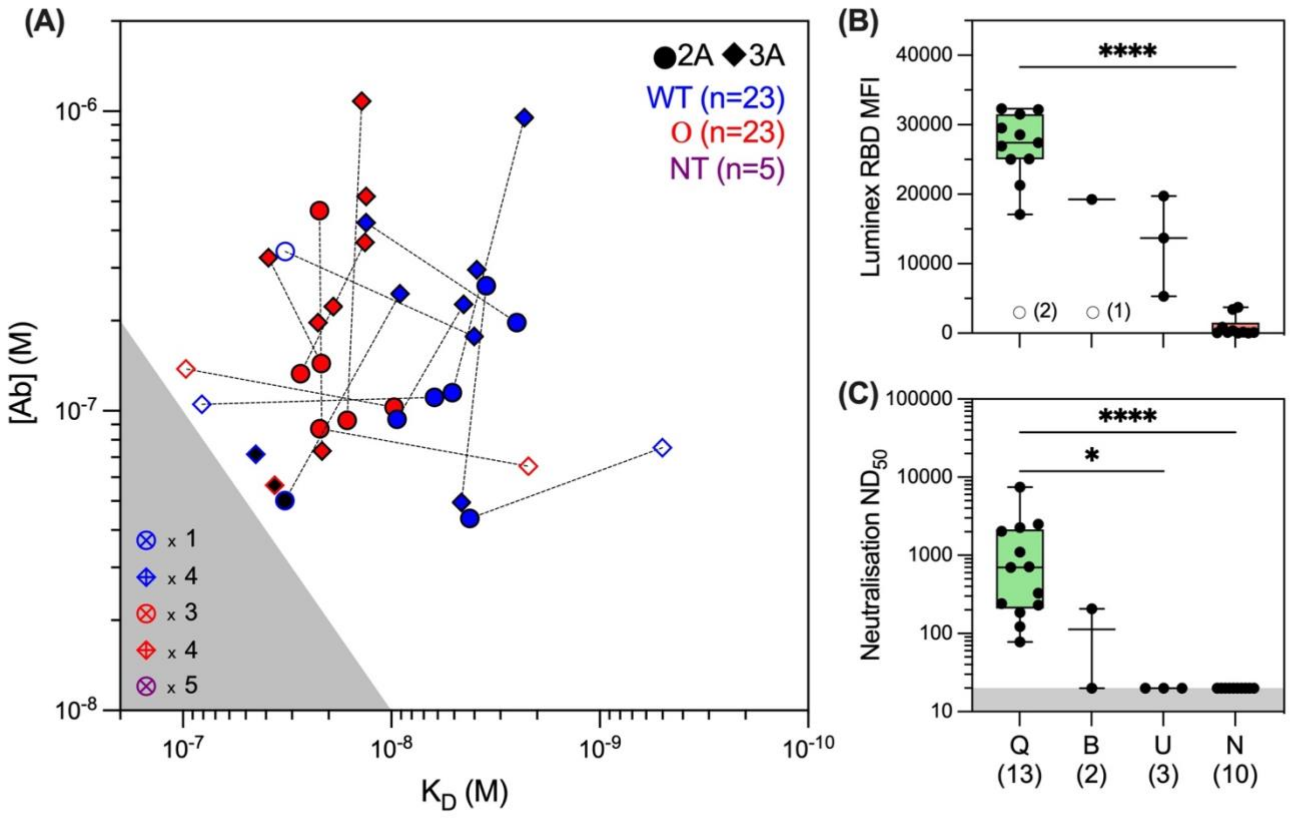


**Supplementary Figure 1. Microfluidic antibody affinity (MAAP), Luminex and neutralisation data for the Rituximab patient cohort. (A)** 2-D scatter plot displaying the best-fit values for antibody binding site affinity (KD, M) and concentration ([Ab], M) measured from MAAP assessment of Rituximab treated vasculitis patient sera taken one month after the 2nd (2A, circles) and 3rd (3A, diamonds) vaccine doses against Wild type (WT, blue symbols) and Omicron (red symbols) Spike RBD domains. Cross-containing symbols represent samples which demonstrated no binding. Unfilled symbols display the measured upper bound 95% confidence intervals for both [Ab] and KD for samples which could not be fully quantified via MAAP due to the inability to constrain the lower bound of the KD value. Black-filled symbols represent samples that were measured at the limit of assay sensitivity (where [Ab]/KD≈1). Purple symbols represent samples which were not assessed via MAAP against either RBD variant, however, are assumed to be non-binders due to their minimal Luminex RBD MFI and negative neutralisation assay data. Grey region represents the unquantifiable range where [Ab]/KD≤1. Dotted lines connect samples from each timepoint from the same patient assessed against the same antigen. **(B)** Categorisation of all samples assessed via MAAP against WT Spike RBD into quantifiable binding (Q=13), quantified but borderline sensitivity (B=2), unquantifiable binding due to inability to constrain the lower bound KD value (U=3), and non-binders (N=10). MAAP quantifiable samples had significantly higher Luminex WT RBD MFI ($p<0.0001$) than non-binders. Unfilled symbols represent samples (number) where Luminex MFI data was unavailable. **(C)** MAAP-quantified samples also demonstrated a significantly higher neutralising capacity (ND50) against WT SARS-CoV-2 than unquantifiable and non-binding samples ($p<0.0001$ and $p=0.0228$, respectively) as measured by viral neutralisation assay. Box blots in B and C represent the median, range and interquartile range for each dataset. Statistical analysis in B and C were carried out via Dunn's multiple comparisons test, presented p-values are two-tailed.

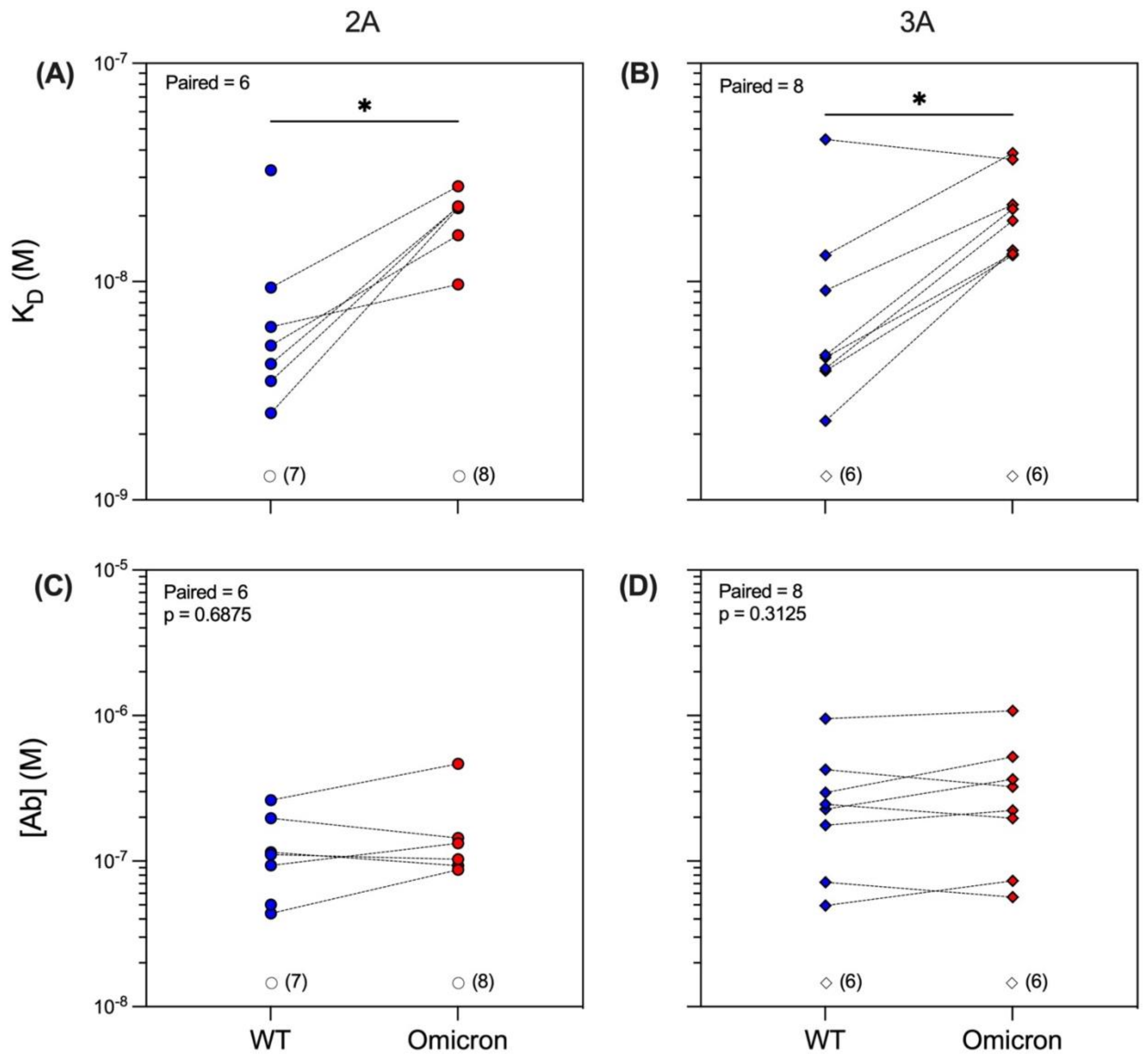


**Supplementary Figure 2. Comparison of anti-SARS-CoV-2 antibody affinity and concentration against the Wild type (WT) and Omicron variants after the second and third vaccine dose in the Rituximab patient cohort.** Comparison of affinity ($K_D$, M) and binding site concentration ([Ab], M) of serum antibodies from Rituximab-treated vasculitis patients showed weaker affinity against the Omicron variant (red symbols) compared to the WT variant (blue symbols). This was consistent in samples taken one month after the 2$^{nd}$ vaccine dose (**A**; 2A, p=0.0312) and after the 3$^{rd}$ vaccine dose (**B**; 3A, p=0.0156). The concentration of antibodies that recognised RBD proteins was found to be similar for WT or Omicron strains at both timepoints (**C**; 2A, p=0.6875, and **D**; 3A, p=0.3125). Dotted lines connect identical samples assessed against different variants. Unfilled symbols represent the samples (number) where MAAP data was unobtainable due to insufficient or unquantifiable binding. P-values presented are two-tailed from the Wilcoxon paired signed-ranks test.

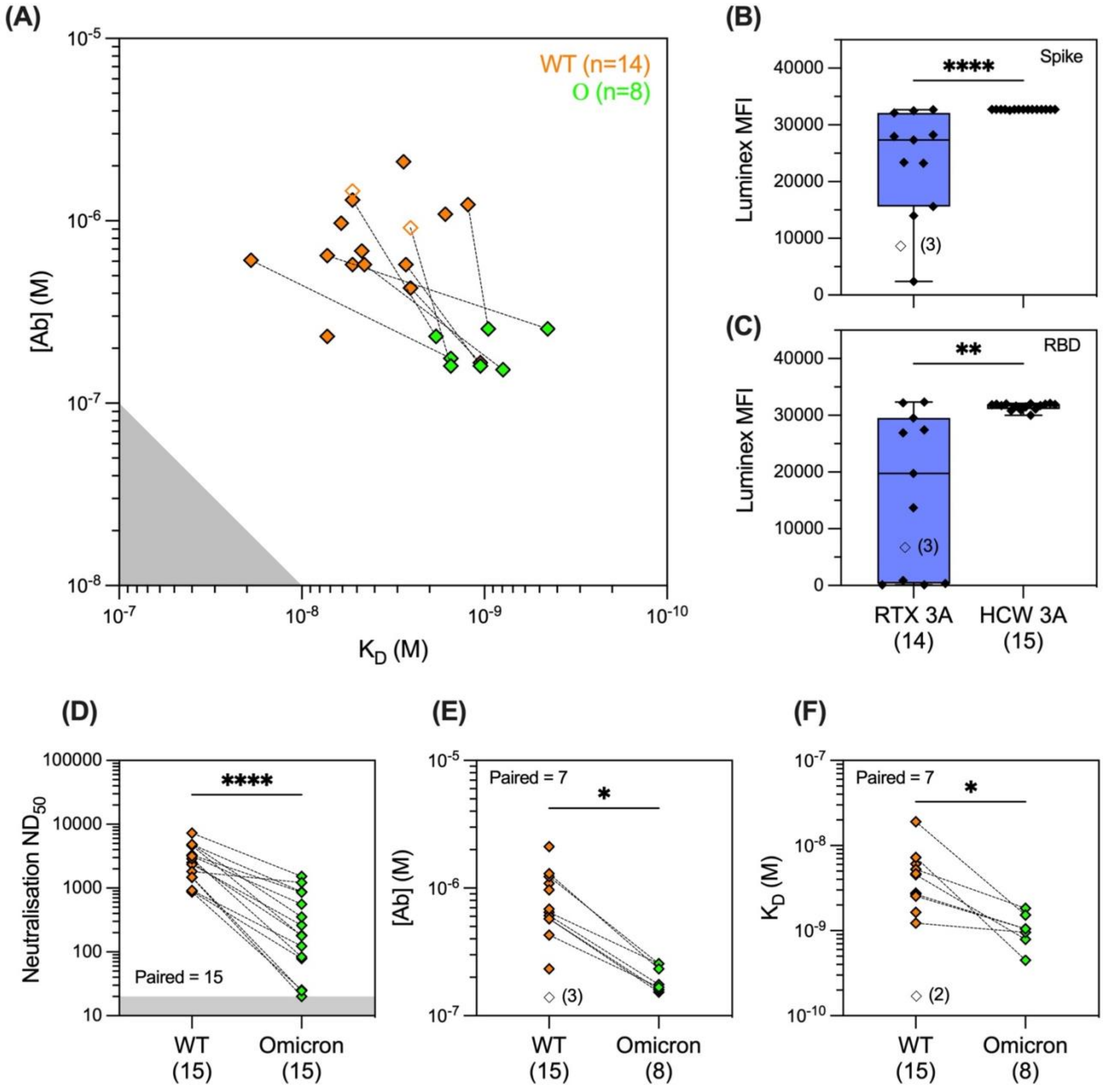


**Supplementary Figure 3. Microfluidic antibody affinity (MAAP), Luminex and neutralisation data for the Health Care Worker (HCW) cohort.** Assessment of healthcare worker sera taken one month after the $3^{rd}$ sensitisation event (3A) using MAAP against WT (orange symbols) and Omicron (green symbols) Spike RBD domains. The 2-D scatter plot maps the best-fit values for antibody affinity ($K_D$, M) and binding site concentration ([Ab], M) **(A)**. Comparison of Luminex anti-Spike (**B**; p<0.0001) and -RBD (**C**; 0.0054) mean fluorescence intensity (MFI) in HCW versus Rituximab treated patients at similar time points (3A: one month after the third exposure). Panel **D** depicts the comparison of neutralisation capacity ($ND_{50}$) in sera from vaccinated healthcare workers at the 3A time point against Omicron versus WT SARS-CoV-22 variant (**D**; p<0.0001). Panel **E** shows the concentration of antibody binding sites that recognised the Omicron variant RBD in comparison to the WT RBD (**E**; p=0.0156) and panel **F** shows the affinity of these antibodies against the two variants (**F**; p=0.0156) at the 3A timepoint. Grey regions represent the assay limits, where in **A** this is the unquantifiable range where [Ab]/$K_D$≤1. In **D** the grey region represents the lower limit of detection for the neutralisation assay. Box blots in **B-C** represent the median, range and interquartile range for each dataset. Unfilled symbols in **A**, **E** and **F** represent samples which could not be fully quantified via MAAP due to the inability to constrain the lower bound of the $K_D$ value. Unfilled symbols in **A** represent samples with unquantifiable binding (U) via MAAP due to inability to constrain the lower bound $K_D$ value, where the displayed values are the measured upper bound 95% confidence interval values for both [Ab] and $K_D$. Unfilled symbols in **B-C** represent the samples (number) that did not have MFI data available. Dotted lines in **A**, **D** and **F** connect the same sample assessed against different variants. P-values presented are two-tailed from the Mann Whitney test for panels **B-C** and Wilcoxon paired signed-ranks test for panels **D-F**. In **D-F**, additional statistical analysis was carried out which compared the ranks of the median via Mann Whitney test, all of which showed a statistically significant difference between the analysed cohorts (**D**; p<0.0001, **E**; p<0.0001, **F**; p=0.0001).

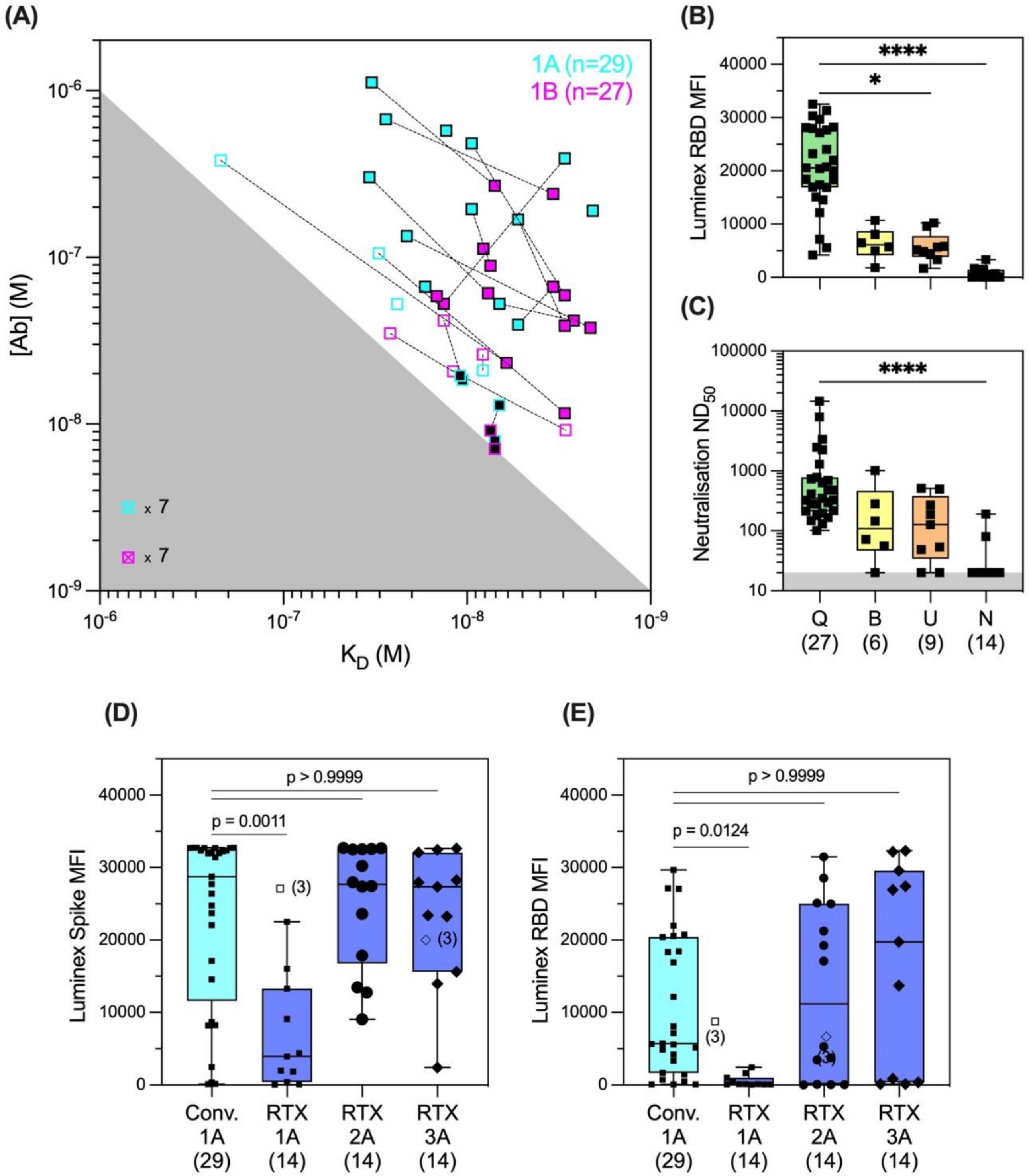


**Supplementary Figure 4. Microfluidic antibody affinity profiling (MAAP), Luminex and neutralisation data for the convalescent cohort. (A)** 2-D scatter plot displaying the best-fit values for antibody affinity ($K_D$, M) and binding site concentration ([Ab], M) measured against Wild type (WT) Spike RBD using MAAP assessment of convalescent patient sera taken one month (1A, teal symbols) and 3 months (1B, magenta symbols) after natural infection during the first wave of SARS-CoV-2. Cross-containing symbols represent samples which demonstrated no binding. Unfilled symbols display the measured upper bound 95% confidence intervals for both [Ab] and $K_D$ for samples which could not be fully quantified via MAAP due to the inability to constrain the lower bound of the $K_D$ value. Black-filled symbols represent samples that were measured at the limit of assay sensitivity (where [Ab]/KD≈1). Grey region represents the unquantifiable range where [Ab]/$K_D$≤1. Dotted lines connect samples taken from the same patient at each timepoint. **(B)** Categorisation of all samples assessed via MAAP against WT Spike RBD into quantifiable binding (Q=27), quantified but borderline sensitivity (B=6), unquantifiable binding due to inability to constrain the lower bound $K_D$ value (U=9), and non-binders (N=14); MAAP quantifiable antibodies had significantly higher Luminex WT RBD MFIs than those in the U and N categories (p=0.0104 and p<0.0001, respectively). **(C)** MAAP quantifiable antibodies also had significantly higher neutralising capacity ($ND_{50}$) than those in the N category (p<0.0001). Grey region represents the lower assay limit. **(D-E)** Comparison of Luminex anti-Spike (**D**) and -RBD (**E**) mean fluorescence intensities (MFI) in the convalescent cohort versus Rituximab-treated (RTX) patients at one month after each sensitisation event. Both anti-Spike and anti-RBD responses were higher in the convalescent versus the Rituximab-treated cohort one month after the first sensitisation event (infection or vaccination, respectively) (Spike; p=0.0011, RBD; p=0.0124), whereas no difference was evident after the second and third vaccination dose in the RTX cohort (all p-values >0.9999). Statistical analyses in **B-E** were carried out via Dunn's multiple comparisons test and presented p-values are two-tailed. Box blots in **B-E** represent the median, range and interquartile range for each dataset. Unfilled symbols represent samples (number) where Luminex MFI data is unavailable.

**(A)**

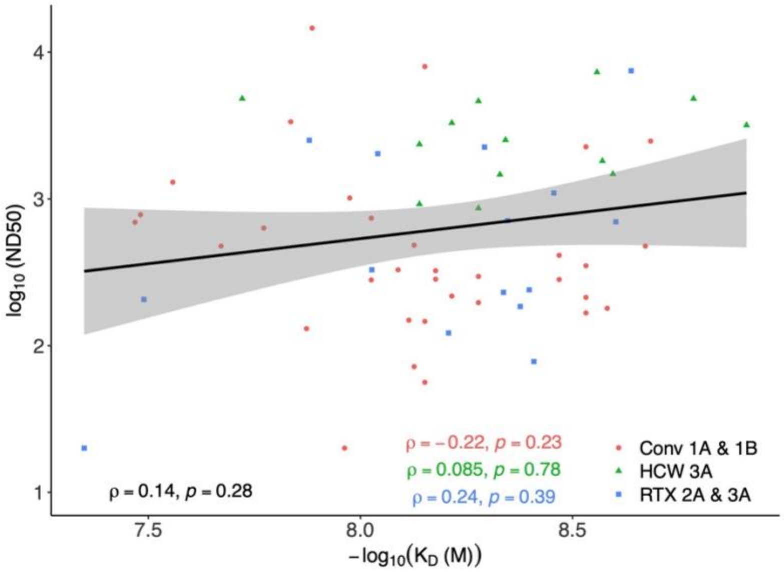
log10 (ND50)
−log10(KD (M))
ρ = 0.14, p = 0.28
ρ = −0.22, p = 0.23
ρ = 0.085, p = 0.78
ρ = 0.24, p = 0.39
Conv 1A & 1B
HCW 3A
RTX 2A & 3A

**(B)**

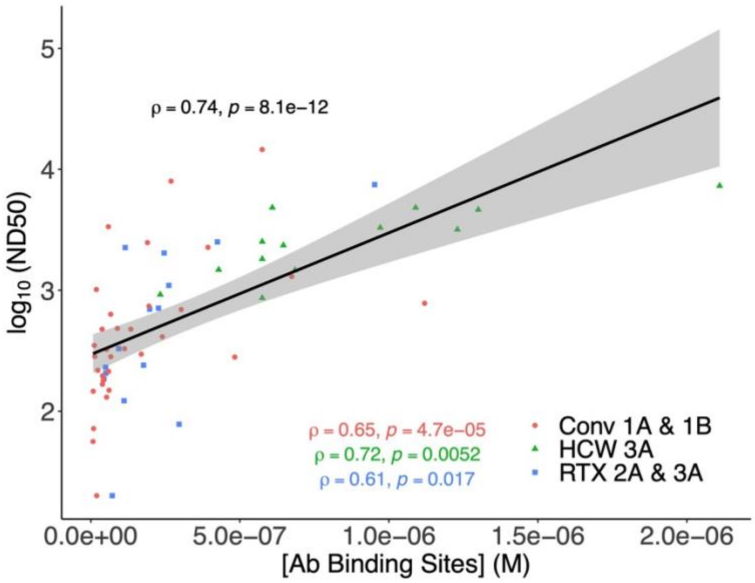
log10 (ND50)
[Ab Binding Sites] (M)
ρ = 0.74, p = 8.1e−12
ρ = 0.65, p = 4.7e−05
ρ = 0.72, p = 0.0052
ρ = 0.61, p = 0.017
Conv 1A & 1B
HCW 3A
RTX 2A & 3A

**(C)**

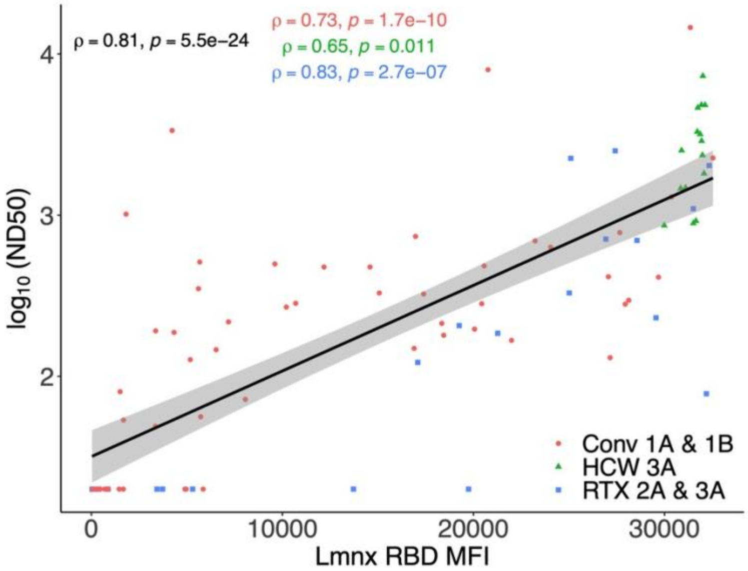
log10 (ND50)
Lmnx RBD MFI
ρ = 0.81, p = 5.5e−24
ρ = 0.73, p = 1.7e−10
ρ = 0.65, p = 0.011
ρ = 0.83, p = 2.7e−07
Conv 1A & 1B
HCW 3A
RTX 2A & 3A

**Supplementary Figure 5. Relationship between serum Wild type SARS-CoV-2 serological read-outs across patient cohorts.** (**A) Relationship between serum Wild type SARS-CoV-2 neutralisation titre (logND$_{50}$) and antibody affinity (-log$_{10}$K$_D$, M) across timepoints and patient cohorts.** Data points from each patient cohort are represented by different shapes. A linear model fit (smooth line) is shown and the Spearman correlation coefficient is depicted, indicating the strength and direction of the correlation between ND$_{50}$ and -log$_{10}$K$_D$. The Spearman correlation coefficients for data in each patient cohort are also shown. (**B) Relationship between serum Wild type SARS-CoV-2 antibody affinity (-log$_{10}$K$_D$, M) and Luminex anti-RBD MFI across timepoints and patient cohorts.** Data points from each patient cohort are represented by different colours. A linear model fit (smooth line) is shown and the Spearman correlation coefficient is depicted, indicating the strength and direction of the correlation between -log$_{10}$K$_D$ and Luminex RBD MFI. The Spearman correlation coefficients for data in each patient cohort are also shown. **(C) Relationship between serum Wild type SARS-CoV-2 antibody binding site concentration ([Ab], M) and Luminex anti-RBD MFI across timepoints and patient cohorts.** Data points from each patient cohort are represented by different colours. A linear model fit (smooth line) is shown and the Spearman correlation coefficient is depicted, indicating the strength and direction of the correlation between [Ab] and Luminex RBD MFI**.** The Spearman correlation coefficients for data in each patient cohort are also shown.

### 2 Supplementary Tables

**Supplementary Table 1. Demographic details of all cohorts studied**. A summary of the available demographic information for each patient cohort assessed within in the study. Clinical data was obtained from electronic medical records and patient interviews and was covered by ethics as described in the methods. *SARS-CoV-2 exposure events for the healthcare worker cohort consisted of two doses of Pfizer vaccine and one natural infection.

| | Vasculitis Patients (n=14) | Healthcare Workers (n=15) one HCW (subjected 213) is missing age/sex/ethnicity. | Convalescent Patients (n=30) |
|---|---|---|---|
| Age (median (range)), years | 66 (24 – 82) | 37 (24 – 60) | 50 (15-85) |
| Sex, male (number (%)) | 8 (57%) | 2 (14%) | 13 (43%) |
| Ethnicity, White British (number (%)) | 13 (93%) | 10 (71%) | 23 (85%) |
| Missing data | | 1 (7%) | |
| Diagnosis | | | |
| ANCA-associated vasculitis | 11 (79%) | - | - |
| SLE | 1 (7%) | - | - |
| IgG4 disease | 1 (7%) | - | - |
| Other* | 1 (7%) | - | - |
| Comorbidities | | | |
| Hypertension | 8 (57%) | - | 8 (27%) |
| Diabetes | 4 (29%) | - | 5 (17%) |
| Chronic lung disease | 4 (29%) | - | 1 (3%) |
| Cardiac disease | 2 (14%) | - | 0 (0%) |
| History of malignancy | 1 (7%) | - | 0 (0%) |
| Vaccines | | | |
| Pfizer x3 doses | 6 (43%) | 15 (100%)* | NA |
| AZ x 3 doses | 1 (7%) | - | NA |
| AZ x2 doses + Pfizer x1 dose | 3 (21%) | - | NA |
| Unknown Combination | 4 (29%) | - | NA |
| GFR (ml/min/m2) (median (IQR)) | 80.5 (63 – 93) | - | - |
| Rituximab | | | |
| Days prior to 1st vaccine dose (median (IQR) | 222.5 (124.5 – 361.25) | - | - |
| Within 6 months | 5 (36%) | - | - |
| Within 12 months | 11 (79%) | - | - |
| Within 5 years | 14 (100%) | - | - |
| Between 1st and 2nd vaccine dose | 3 (21%) | - | - |
| Between 2nd and 3rd vaccine dose | 6 (43%) | - | - |
| Cyclophosphamide | | | |
| Between 1st and 2nd vaccine dose | 2 (14%) | - | - |
| Between 2nd and 3rd vaccine dose | 1 (7%) | - | - |
| Time between vaccines (median (range)), days | | | |
| Time between vaccines (median (range)), days | | | |
| 1st and 2nd vaccines | 77 (58 – 119) | missing | - |
| 2nd and 3rd vaccines | 190 (147 – 201) | missing | - |

**Supplementary Table 2. All data for the Rituximab-treated vasculitis patient cohort.** A summary of the samples obtained from each patient within the rituximab-treated vasculitis cohort (ethics reference: 20/EM/180) and all the raw data gathered from Luminex (RBS spike, Spike S1 and Nucleocapsid), pseudotype neutralisation (Neut. $ND_{50}$), and microfluidic antibody affinity profiling (MAAP) assays for each sample. 2A and 3A represent samples taken at one month post second and third vaccine dose, respectively. Q/B/U/N represents whether the sample assessed via MAAP could be fully quantified (Q), whether it was near the border of detectible sensitivity (B; [Ab]/KD<2), whether it was unquantifiable due to the inability to effectively constrain the lower 95% confidence interval (U), or whether there was no binding observed (N). * represents samples which were not assessed via MAAP but are assumed to be non-binders due to minimal Luminex signal and negative neutralisation. [X] represents values that may be raised due to the $[Ab]/K_D<2$. [Y] represents values which may be inaccurate due to the lack of lower 95% $K_D$ constraints. – means that data was not available for sample using this assay.

| Patient | Sample Timepoint | RBD Variant | Luminex (MFI) | | | Neut. ND50 | MAAP Data | | | | | | | |
|---|---|---|---|---|---|---|---|---|---|---|---|---|---|---|
| | | | | | | | Q/B/U/N | $K_D$ (nM) | | | [Ab] (nM) | | | [Ab]/KD |
| | | | RBD | Spike | N | | | Best Fit | Upper | Lower | Best Fit | Upper | Lower | |
| 101 | 2A | WT | 25094.5 | 27406.0 | 972.5 | 2254.0 | Q | 5.1 | 8.5 | 2.4 | 115.2 | 137.5 | 87.2 | 22.59 |
| | | Omicron | - | - | - | 125.5 | Q | 16.3 | 22.3 | 10.5 | 93.0 | 122.2 | 68.1 | 5.71 |
| | 3A | WT | - | - | - | 7468.0 | Q | 2.3 | 10.3 | 1.2 | 952.1 | 2663 | 702.3 | 413.96 |
| | | Omicron | - | - | - | 648.7 | Q | 13.9 | 27.4 | 6.7 | 1082.3 | 1960.6 | 749.5 | 77.86 |
| 102 | 2A | WT | 33.0 | 17847.5 | 78.5 | <20 | N* | - | - | - | - | - | - | - |
| | | Omicron | - | - | - | <20 | N* | - | - | - | - | - | - | - |
| | 3A | WT | 369.5 | 15629.0 | 99.5 | <20 | N | - | - | - | - | - | - | - |
| | | Omicron | - | - | - | <20 | N | - | - | - | - | - | - | - |
| 103 | 2A | WT | 52.8 | 23606.0 | 63.3 | <20 | N* | - | - | - | - | - | - | - |
| | | Omicron | - | - | - | <20 | N* | - | - | - | - | - | - | - |
| | 3A | WT | 874.5 | 23405.0 | 68.0 | <20 | N | - | - | - | - | - | - | - |
| | | Omicron | - | - | - | <20 | N | - | - | - | - | - | - | - |
| 104 | 2A | WT | 28556.8 | 32514.0 | 248.0 | 697.7 | Q | 2.5 | 3.4 | 1.9 | 197.3 | 246.2 | 173.6 | 78.92 |
| | | Omicron | - | - | - | 49.0 | Q | 21.7 | 35.5 | 12.5 | 143.9 | 204.3 | 104.0 | 6.63 |
| | 3A | WT | 27415.0 | 27961.0 | 879.0 | 2509.0 | Q | 13.2 | 25.1 | 6.6 | 424.6 | 613.5 | 260.9 | 32.17 |
| | | Omicron | - | - | - | 314.2 | Q | 38.8 | 148.1 | 11.4 | 324.5 | 1076.0 | 135.8 | 8.36 |
| 105 | 2A | WT | 21268.0 | 32678.0 | 1249.0 | 185.0 | Q | 4.2 | 6.6 | 2.6 | 43.7 | 60.8 | 35.8 | 10.40 |
| | | Omicron | - | - | - | <20 | Q | 22.0 | 30.6 | 14.5 | 87.2 | 119.3 | 62.5 | 3.96 |
| | 3A | WT | 19741.0 | 28266.5 | 138.0 | <20 | U | 0.0[y] | 0.5[y] | 0.0[y] | 54.8[y] | 75.4[y] | 6.8[y] | -[y] |
| | | Omicron | - | - | - | <20 | U | 7.1[y] | 2.2[y] | 0.0[y] | 9.8[y] | 65.4[y] | 4.1[y] | 1.38[y] |
| 106 | 2A | WT | 3730.0 | 30226.0 | 282.0 | <20 | N | - | - | - | - | - | - | - |
| | | Omicron | - | - | - | <20 | N | - | - | - | - | - | - | - |
| | 3A | WT | 32188.0 | 32484.0 | 325.5 | 78.0 | Q | 3.9 | 5.0 | 3.0 | 296.2 | 329.7 | 263.5 | 75.95 |
| | | Omicron | - | - | - | 153.3 | Q | 13.2 | 24.3 | 6.1 | 519.7 | 718.8 | 371.3 | 39.37 |
| 107 | 2A | WT | 42.8 | 9046.5 | 496.8 | <20 | N* | - | - | - | - | - | - | - |
| | | Omicron | - | - | - | <20 | N* | - | - | - | - | - | - | - |
| | 3A | WT | 127.0 | 2398.0 | 307.0 | <20 | N | - | - | - | - | - | - | - |
| | | Omicron | - | - | - | <20 | N | - | - | - | - | - | - | - |
| 108 | 2A | WT | 5301.0 | 27451.0 | 407.0 | <20 | U | 0.2[y] | 32.3[y] | 0.0[y] | 205.9[y] | 340.7[y] | 102.2[y] | 1029.50[y] |
| | | Omicron | - | - | - | <20 | N | - | - | - | - | - | - | - |
| | 3A | WT | - | - | - | 240.3 | Q | 4.0 | 6.1 | 2.7 | 176.8 | 222.4 | 135.7 | 44.20 |
| | | Omicron | - | - | - | <20 | Q | 19.0 | 43.3 | 8.6 | 223.1 | 383.1 | 111.5 | 11.74 |
| 109 | 2A | WT | 25015.3 | 32554.8 | 48.8 | 329.0 | Q | 9.4 | 12.7 | 6.3 | 93.7 | 117.1 | 71.9 | 9.97 |
| | | Omicron | - | - | - | <20 | Q | 27.3 | 39.1 | 18.1 | 132.8 | 194.8 | 99.7 | 4.86 |
| | 3A | WT | 26930.5 | 27356.0 | 99.0 | 711.0 | Q | 4.5 | 5.6 | 3.3 | 227.1 | 254.9 | 197.0 | 50.47 |
| | | Omicron | - | - | - | <20 | Q | 13.4 | 17.3 | 10.2 | 365.3 | 423.6 | 313.1 | 27.26 |

| | | | | | | | | | | | | | | |
|---|---|---|---|---|---|---|---|---|---|---|---|---|---|---|
| 110 | 2A | WT | 19252.5 | 27957.3 | 6181.0 | 206.5 | B | 32.4[x] | 68.8[x] | 7.9[x] | 50.2[x] | 117.2[x] | 17.5[x] | 1.55[x] |
| | | Omicron | - | - | - | <20 | N | - | - | - | - | - | - | - |
| | 3A | WT | 32331.0 | 32661.5 | 6564.0 | 2033.0 | Q | 9.1 | 10.8 | 7.3 | 245.9 | 271.4 | 218.3 | 27.02 |
| | | Omicron | - | - | - | 177.6 | Q | 22.5 | 33.3 | 15.4 | 197.3 | 257.7 | 145.4 | 8.77 |
| 111 | 2A | WT | 103.5 | 12767.5 | 172.0 | <20 | N* | - | - | - | - | - | - | - |
| | | Omicron | - | - | - | <20 | N* | - | - | - | - | - | - | - |
| | 3A | WT | 135.5 | 13983.5 | 202.5 | <20 | N | - | - | - | - | - | - | - |
| | | Omicron | - | - | - | <20 | N | - | - | - | - | - | - | - |
| 112 | 2A | WT | 31501.8 | 32716.3 | 704.8 | 1097.0 | Q | 3.5 | 5.3 | 2.2 | 261.7 | 315.9 | 217.2 | 74.77 |
| | | Omicron | - | - | - | 61.4 | Q | 22.1 | 33.7 | 13.8 | 466.2 | 584.4 | 367.9 | 21.10 |
| | 3A | WT | 29550.5 | 32095.0 | 1574.0 | 230.9 | Q | 4.6 | 9.4 | 1.5 | 49.5 | 73.6 | 29.9 | 10.76 |
| | | Omicron | - | - | - | <20 | Q | 21.5 | 34.2 | 13.0 | 73.4 | 109.1 | 46.5 | 3.41 |
| 113 | 2A | WT | 3439.7 | 13478.3 | 79.0 | <20 | N* | - | - | - | - | - | - | - |
| | | Omicron | - | - | - | <20 | N* | - | - | - | - | - | - | - |
| | 3A | WT | - | - | - | <20 | B | 44.8[x] | 81.6[x] | 17.0[x] | 71.7[x] | 159.5[x] | 32.2[x] | 1.60[x] |
| | | Omicron | - | - | - | <20 | B | 36.3[x] | 66.9[x] | 13.4[x] | 56.5[x] | 133.4[x] | 26.6[x] | 1.56[x] |
| 114 | 2A | WT | 17083.5 | 32590.0 | 1504.0 | 122.0 | Q | 6.2 | 10.9 | 3.4 | 111.3 | 146.9 | 80.4 | 17.95 |
| | | Omicron | - | - | - | <20 | Q | 9.7 | 13.3 | 7.2 | 102.9 | 118.4 | 88.7 | 10.61 |
| | 3A | WT | 13717.5 | 23244.0 | 1078.0 | <20 | U | 8.2[y] | 81.0[y] | 0.0[y] | 9.1[y] | 105.1[y] | 0.0[y] | 1.11[y] |
| | | Omicron | - | - | - | <20 | U | 1.0[y] | 96.9[y] | 0.0[y] | 12.9[y] | 138.2[y] | 3.7[y] | 12.90[y] |

**Supplementary Table 3. All data for the healthcare worker cohort.** A summary of the samples obtained from each patient within the rituximab-treated vasculitis cohort (ethics reference: 17/EE/0025) and all the raw data gathered from Luminex (RBS spike, Spike S1 and Nucleocapsid), pseudotype neutralisation (Neut. $ND_{50}$), and microfluidic antibody affinity profiling (MAAP) assays for each sample. 3A represents samples taken at one month post third SARS-CoV-2 exposure (2 vaccines and one natural infection). Q/B/U/N represents whether the sample assessed via MAAP could be fully quantified (Q), whether it was near the border of detectible sensitivity (B; [Ab]/KD between 1-2), whether it was unquantifiable due to the inability to effectively constrain the lower 95% confidence interval (U), or whether there was no binding observed (N). [Y] represents values which may be inaccurate due to the lack of lower 95% $K_D$ constraints. – means that data was not available for sample using this assay.

| Patient | Sample Timepoint | RBD Variant | Luminex (MFI) | | | Neut. ND50 | MAAP Data | | | | | | | |
|---|---|---|---|---|---|---|---|---|---|---|---|---|---|---|
| | | | | | | | Q/B/U/N | $K_D$ (nM) | | | [Ab] (nM) | | | [Ab]/KD |
| | | | RBD | Spike | N | | | Best Fit | Upper | Lower | Best Fit | Upper | Lower | |
| 201 | 3A | WT | 32121.0 | 32728.5 | 22051.0 | 4816 | Q | 1.6 | 3.4 | 0.3 | 1090.0 | 1460.0 | 483.0 | 664.63 |
| | | Omicron | - | - | - | 873.9 | - | - | - | - | - | - | - | - |
| 202 | 3A | WT | 31714.0 | 32711.0 | 4931.0 | 3293 | Q | 6.1 | 10.9 | 1.2 | 971.0 | 1550.0 | 429.0 | 159.44 |
| | | Omicron | - | - | - | 857.8 | - | - | - | - | - | - | - | - |
| 203 | 3A | WT | 30891.0 | 32735.0 | 6116.0 | 2522 | Q | 4.6 | 8.2 | 2.2 | 575.0 | 815.0 | 382.0 | 126.37 |
| | | Omicron | - | - | - | 182.7 | Q | 0.8 | 1.8 | 0.1 | 153.0 | 222.0 | 86.9 | 193.43 |
| 204 | 3A | WT | 31984.0 | 32737.0 | 11615.0 | 2355 | Q | 7.3 | 10.9 | 4.1 | 646.0 | 815.0 | 483.0 | 88.98 |
| | | Omicron | - | - | - | 260.3 | Q | 0.5 | 1.2 | 0.1 | 256.0 | 373.0 | 184.0 | 568.89 |
| 205 | 3A | WT | 31505.0 | 32735.0 | 412.0 | 891.7 | U | 2.5[y] | 5.3[y] | 0.0[y] | 971.0[y] | 1460.0[y] | 340.0[y] | 382.28[y] |
| | | Omicron | - | - | - | 79.01 | - | - | - | - | - | - | - | - |
| 206 | 3A | WT | 31939.0 | 32735.0 | 5814.5 | 2889 | U | 0.0[y] | 2.5[y] | 0.0[y] | 609.0[y] | 916.0[y] | 119.0[y] | -[y] |
| | | Omicron | - | - | - | 83.09 | Q | 1.5 | 3.6 | 0.7 | 160.0 | 233.0 | 121.0 | 104.58 |
| 207 | 3A | WT | 31862.0 | 32735.0 | 20377.0 | 3178 | Q | 1.2 | 2.2 | 0.1 | 1230.0 | 1640.0 | 646.0 | 1000.00 |
| | | Omicron | - | - | - | 562.2 | Q | 1.0 | 1.8 | 0.3 | 256.0 | 324.0 | 176.0 | 268.34 |
| 208 | 3A | WT | 31740.0 | 32737.0 | 5096.0 | 4639 | Q | 5.3 | 9.4 | 1.5 | 1300.0 | 1950.0 | 609.0 | 246.68 |
| | | Omicron | - | - | - | 178.9 | Q | 1.8 | 4.3 | 0.5 | 233.0 | 295.0 | 176.0 | 126.63 |
| 209 | 3A | WT | 31929.0 | 32734.0 | 4667.0 | 4824 | Q | 19.0 | 38.2 | 7.5 | 609.0 | 971.0 | 382.0 | 32.05 |
| | | Omicron | - | - | - | 353.8 | Q | 1.5 | 2.7 | 0.7 | 176.0 | 233.0 | 126.0 | 115.03 |
| 210 | 3A | WT | 29985.0 | 32713.5 | 8161.0 | 864 | Q | 5.3 | 10.6 | 2.1 | 575.0 | 1030.0 | 361.0 | 109.11 |
| | | Omicron | - | - | - | 25.5 | - | - | - | - | - | - | - | - |
| 211 | 3A | WT | 30841.0 | 32603.5 | 5122.0 | 1466 | Q | 4.7 | 9.4 | 1.6 | 685.0 | 1030.0 | 405.0 | 146.06 |
| | | Omicron | - | - | - | <20 | - | - | - | - | - | - | - | - |
| 212 | 3A | WT | 32004.0 | 32734.0 | 27589.0 | 7297 | Q | 2.8 | 5.3 | 0.9 | 2110.0 | 2760.0 | 1360.0 | 761.73 |
| | | Omicron | - | - | - | 1536 | - | - | - | - | - | - | - | - |
| 213 | 3A | WT | 32068.0 | 32735.0 | 23521.0 | 1814 | Q | 2.7 | 5.9 | 0.7 | 575.0 | 916.0 | 405.0 | 213.75 |
| | | Omicron | - | - | - | 1225 | Q | 1.1 | 3.0 | 0.4 | 160.0 | 309.0 | 115.0 | 152.38 |
| 214 | 3A | WT | 31102.0 | 32731.0 | 17863.0 | 1478 | Q | 2.5 | 3.9 | 1.1 | 429.0 | 542.0 | 286.0 | 168.90 |
| | | Omicron | - | - | - | 24.6 | Q | 1.1 | 2.0 | 0.3 | 167.0 | 223.0 | 109.0 | 159.05 |
| 215 | 3A | WT | 31643.0 | 32735.0 | 1087.0 | 921.3 | Q | 7.3 | 13.4 | 3.3 | 233.0 | 330.0 | 155.0 | 32.09 |
| | | Omicron | - | - | - | 123.3 | - | - | - | - | - | - | - | - |

**Supplementary Table 4. All data for the convalescent patient cohort.** A summary of the samples obtained from each patient within the convalescent patient cohort who were exposed to ancestral strain of SARS-CoV-2 during the first wave (ethics reference: 17/EE/0025) and all the raw data gathered from Luminex (RBS spike, Spike S1 and Nucleocapsid), pseudotype neutralisation (Neut. $ND_{50}$), and microfluidic antibody affinity profiling (MAAP) assays for each sample. 1A and 1B represents samples taken at one- and three-months after the first positive COVID test, respectively. Q/B/U/N represents whether the sample assessed via MAAP could be fully quantified (Q), whether it was near the border of detectible sensitivity (B; [Ab]/KD between 1-2), whether it was unquantifiable due to the inability to effectively constrain the lower 95% confidence interval (U), or whether there was no binding observed (N). [X] represents values that may be raised due to the [Ab]/$K_D$<2. [Y] represents values which may be inaccurate due to the lack of lower 95% $K_D$ constraints. – means that data was not available for sample using this assay.

| Patient | Sample Timepoint | Variant | Luminex (MFI) | | | Neut. ND50 | MAAP Data | | | | | | | |
|---|---|---|---|---|---|---|---|---|---|---|---|---|---|---|
| | | | | | | | Q/B/U/N | $K_D$ (nM) | | | [Ab] (nM) | | | [Ab]/KD |
| | | | RBD | Spike | N | | | Best Fit | Upper | Lower | Best Fit | Upper | Lower | |
| 301 | 1A | WT | 28211.5 | 32721.0 | 31937.0 | 2478.0 | Q | 2.07 | 5.92 | 0.455 | 190 | 303 | 106 | 91.79 |
| 302 | 1A | WT | 3365.5 | 24755.0 | 31038.5 | 191.6 | N | - | - | - | - | - | - | - |
| | 1B | | 4228.0 | 25393.5 | 31776.0 | 3354.0 | Q | 14.6 | 40.5 | 2.94 | 58.5 | 99.1 | 17.9 | 4.01 |
| 303 | 1A | WT | 27941.0 | 32733.0 | 32123.0 | 280.4 | Q | 9.4 | 34.0 | 1.2 | 483.0 | 1090.0 | 190.0 | 51.22 |
| | 1B | | 18341.0 | 31937.0 | 31068.0 | 213.2 | Q | 2.9 | 5.3 | 1.2 | 59.2 | 94.3 | 41.7 | 20.14 |
| 304 | 1A | WT | 31347.0 | 32737.0 | 32735.5 | 14580.0 | Q | 13.0 | 23.3 | 7.7 | 575.0 | 916.0 | 429.0 | 44.23 |
| 305 | 1A | WT | 9607.0 | 27754.5 | 31715.0 | 498.2 | U | 8.4[y] | 30.3[y] | 0.0[y] | 29.4[y] | 106.0[y] | 5.1[y] | 3.50[y] |
| | 1B | | 5607.0 | 21572.5 | 16095.0 | 350.2 | Q | 2.9 | 14.6 | 0.2 | 11.6 | 37.1 | 4.1 | 3.95 |
| 306 | 1A | WT | 890.0 | 8678.0 | 6978.0 | <20 | N | - | - | - | - | - | - | - |
| | 1B | | 702.0 | 6260.0 | 3021.0 | <20 | N | - | - | - | - | - | - | - |
| 307 | 1A | WT | 1507.0 | 8239.5 | 20981.0 | 80.37 | N | - | - | - | - | - | - | - |
| | 1B | | 1673.0 | 12215.0 | 14433.0 | <20 | N | - | - | - | - | - | - | - |
| 308 | 1A | WT | 4329.0 | 28763.0 | 28407.0 | 187.4 | U | 3.9[y] | 8.2[y] | 0.0[y] | 9.6[y] | 21.0[y] | 2.9[y] | 2.44[y] |
| | 1B | | 5179.0 | 27945.5 | 20467.0 | 127.0 | U | 1.9[y] | 8.2[y] | 0.0[y] | 8.2[y] | 26.2[y] | 1.8[y] | 4.29[y] |
| 309 | 1A | WT | 89.0 | 116.0 | 282.0 | <20 | N | - | - | - | - | - | - | - |
| | 1B | | 82.0 | 74.0 | 203.5 | <20 | N | - | - | - | - | - | - | - |
| 310 | 1A | WT | 4966.0 | 14558.0 | 31916.5 | <20 | B | 10.9[x] | 46.9[x] | 0.2[x] | 19.5[x] | 97.1[x] | 6.1[x] | 1.79[x] |
| | 1B | | 4899.5 | 13056.0 | 30355.0 | <20 | U | 8.15[y] | 26.2[y] | 0.0[y] | 7.92[y] | 35.0[y] | 1.16[y] | 0.97[y] |
| 311 | 1A | WT | 1683.0 | 8207.0 | 14663.0 | 53.6 | U | 8.4[y] | 24.0[y] | 0.0[y] | 14.2[y] | 52.6[y] | 2.3[y] | 1.69[y] |
| | 1B | | 1459.5 | 8346.0 | 10403.0 | <20 | N | - | - | - | - | - | - | - |
| 312 | 1A | WT | 23224.0 | 32375.5 | 32074.5 | 693.0 | Q | 34.0 | 86.4 | 7.5 | 303.0 | 864.0 | 164.0 | 8.91 |
| | 1B | | 16894.0 | 31319.0 | 31706.0 | 149.1 | Q | 7.7 | 14.6 | 2.7 | 60.9 | 116.0 | 38.2 | 7.92 |
| 313 | 1A | WT | 27659.0 | 32732.5 | 32737.0 | 780.3 | Q | 33.0 | 150.0 | 8.2 | 1120.0 | 1870.0 | 303.0 | 33.94 |
| | 1B | | 20759.0 | 32123.0 | 31280.5 | 7983 | Q | 7.1 | 30.3 | 0.5 | 269.0 | 609.0 | 59.2 | 38.16 |
| 314 | 1A | WT | 16968.0 | 32278.0 | 32428.5 | 738.9 | Q | 9.4 | 21.3 | 2.3 | 195.0 | 330.0 | 81.5 | 20.68 |
| | 1B | | 20561.5 | 32018.5 | 31996.0 | 484.2 | Q | 7.5 | 15.0 | 2.1 | 89.0 | 164.0 | 44.2 | 11.91 |
| 315 | 1A | WT | 32532.0 | 32734.5 | 32736.0 | 2264.0 | Q | 2.9 | 11.9 | 1.2 | 393.0 | 1340.0 | 277.0 | 133.67 |
| | 1B | | 27150.0 | 31419.0 | 31279.0 | 130.6 | Q | 13.4 | 24.0 | 3.7 | 52.7 | 97.1 | 26.2 | 3.93 |
| 316 | 1A | WT | 10206.0 | 26395.0 | 30329.0 | 269.1 | U | 13.4[y] | 220.0[y] | 0.0[y] | 20.7[y] | 382.0[y] | 6.5[y] | 1.54[y] |
| | 1B | | 7177.0 | 22897.0 | 19629.5 | 217.8 | Q | 6.1 | 16.9 | 0.6 | 23.3 | 46.9 | 6.5 | 3.83 |
| 317 | 1A | WT | 84.0 | 366.0 | 114.0 | <20 | N | - | - | - | - | - | - | - |
| | 1B | | 85.0 | 413.0 | 139.5 | <20 | N | - | - | - | - | - | - | - |

| | | | | | | | | | | | | | | |
|---|---|---|---|---|---|---|---|---|---|---|---|---|---|---|
| 318 | 1A | WT | 1817.5 | 17131.5 | 31787.0 | 1015.0 | B | 10.6[x] | 38.2[x] | 0.7[x] | 18.4[x] | 94.3[x] | 7.3[x] | 1.74[x] |
| | 1B | | 5674.0 | 26432.0 | 28345.0 | 512.2 | U | 4.7[y] | 13.4[y] | 0.0[y] | 11.6[y] | 41.7[y] | 4.1[y] | 2.47[y] |
| 319 | 1A | WT | 291.0 | 147.5 | 336.5 | <20 | N | - | - | - | - | - | - | - |
| | 1B | | 464.0 | 259.0 | 969.5 | <20 | N | - | - | - | - | - | - | - |
| 320 | 1A | WT | 5851.0 | 23703.0 | 11289.0 | <20 | U | 4.7[y] | 11.9[y] | 0.0[y] | 5.8[y] | 20.7[y] | 2.0[y] | 1.23[y] |
| | 1B | | 3355.0 | 13418.5 | 4301.0 | 49.0 | U | 1.4[y] | 2.9[y] | 0.0[y] | 4.1[y] | 9.2[y] | 1.8[y] | 2.85[y] |
| 321 | 1A | WT | 28124.5 | 32728.0 | 31844.0 | 296.3 | Q | 5.3 | 10.9 | 0.3 | 169.0 | 269.0 | 52.7 | 32.07 |
| | 1B | | 21992.0 | 32163.5 | 26469.0 | 167.2 | Q | 2.9 | 5.3 | 0.9 | 38.8 | 69.4 | 29.0 | 13.20 |
| 322 | 1A | WT | 30362.0 | 32734.5 | 32735.0 | 1300.0 | Q | 27.7 | 103.0 | 6.3 | 674.0 | 1500.0 | 281.0 | 24.33 |
| | 1B | | 29678.0 | 32470.0 | 31842.0 | 412.6 | Q | 3.4 | 10.9 | 0.1 | 240.0 | 382.0 | 134.0 | 70.59 |
| 323 | 1B | WT | 15070.5 | 31792.5 | 32546.0 | 328.8 | Q | 8.2 | 16.9 | 1.9 | 113.0 | 192.0 | 44.9 | 13.87 |
| 324 | 1A | WT | 20056.0 | 32369.0 | 31792.5 | 196.4 | Q | 5.3 | 10.9 | 1.9 | 39.4 | 58.5 | 20.4 | 7.48 |
| | 1B | | 20432.0 | 31670.5 | 28798.0 | 282.3 | Q | 3.4 | 10.9 | 0.8 | 66.7 | 113.0 | 34.5 | 19.62 |
| 325 | 1A | WT | 10694.0 | 31967.0 | 31073.5 | 283.8 | B | 6.7[x] | 11.9[x] | 0.5[x] | 13.0[x] | 29.4[x] | 5.8[x] | 1.95[x] |
| | 1B | | 8062.0 | 30697.0 | 21154.0 | 71.9 | B | 7.5[x] | 19.0[x] | 1.2[x] | 9.2[x] | 33.0[x] | 4.6[x] | 1.23[x] |
| 326 | 1A | WT | 14585.0 | 31423.5 | 32736.0 | 476.8 | Q | 21.3 | 5.6 | 0.4 | 134.0 | 67.0 | 23.8 | 6.29 |
| | 1B | | 12174.0 | 30130.0 | 31422.0 | 101.5 | Q | 2.1 | 42.9 | 5.3 | 37.7 | 269.0 | 46.9 | 17.70 |
| 327 | 1A | WT | 77.0 | 2470.0 | 553.0 | <20 | N | - | - | - | - | - | - | - |
| | 1B | | 77.0 | 2483.0 | 457.0 | <20 | N | - | - | - | - | - | - | - |
| 328 | 1A | WT | 17395.0 | 31999.5 | 31133.0 | 324.2 | Q | 6.7 | 11.9 | 2.9 | 52.7 | 94.3 | 33.0 | 7.92 |
| | 1B | | 18443.0 | 31608.0 | 26901.0 | 179.9 | Q | 2.6 | 6.7 | 0.6 | 41.7 | 66.5 | 18.4 | 15.92 |
| 329 | 1A | WT | 24032.0 | 32023.0 | 32023.0 | 633.0 | Q | 16.9 | 34.0 | 5.3 | 66.5 | 169.0 | 37.1 | 3.93 |
| 330 | 1A | WT | 6530.0 | 22076.0 | 22076.0 | 146.3 | B | 7.1[x] | 16.9* | 0.7[x] | 7.9[x] | 109.0[x] | 0.9[x] | 1.12[x] |
| | 1B | | 5723.5 | 19402.0 | 24526.5 | 56.2 | B | 7.1[x] | 16.9* | 0.7[x] | 7.1[x] | 23.3[x] | 2.2[x] | 1.01[x] |
